# Entanglement-governed protein networks enable mechanically adaptive artificial skin for transplantation-scale skin replacement

**DOI:** 10.64898/2026.06.11.731551

**Authors:** Liping Wang, Yinan Sun, Xing Liu, Ruoxuan Wang, Jinxia Huang, Wenbo Wang, Kongxi Fan, Jia Bai, Zhiying Dong, Shuang Jia, Yan Xia, Shubin Li, Liyao Wang, Yuhao Chen, Yitian Du, Xinyu Li

## Abstract

Artificial skin substitutes that simultaneously achieve mechanical robustness, regenerative bioactivity, and transplantation-scale tissue integration remain challenging to engineer. Here we report a mechanically adaptive bilayer artificial skin based on entanglement-mediated protein networks. By integrating protein chain entanglement, flexible molecular linkers, and photo-triggered intermolecular crosslinking, we establish a hierarchically organized protein matrix with enhanced toughness, structural adaptability, and regenerative compatibility. Spatial biofunctionalization further enables integration of an antibacterial Zn²⁺-coordinated epidermal layer and a regenerative CLP–EGF-functionalized dermal layer within a unified construct. The engineered skin promotes cellular proliferation through PI3K–AKT–mTOR activation, exhibits sustained antibacterial activity, and supports large-area full-thickness skin replacement covering approximately 40% of the dorsal skin surface in mice. The construct further accelerates diabetic wound repair and extracellular matrix remodeling in vivo. These findings establish entanglement-mediated protein engineering as a strategy for mechanically adaptive regenerative biomaterials and provide a platform for transplantation-scale skin regeneration.

## Introduction

Skin defects, particularly severe burn injuries, chronic pathological wounds, and large-area full-thickness tissue loss, remain a major clinical challenge worldwide. Skin substitution has emerged as one of the most promising therapeutic strategies for the treatment of extensive and non-healing wounds in reconstructive medicine^[1, 2]^. However, despite decades of development, clinically available skin substitutes still face substantial limitations in terms of bio-integration, mechanical stability, and long-term functional regeneration.

Autologous and allogeneic skin grafts remain the primary clinical approaches for skin transplantation^[3, 4]^. Although autologous grafting is considered the clinical gold standard owing to its superior histocompatibility, its application is severely restricted by limited donor-site availability and secondary tissue injury, particularly in patients with extensive burns or repeated grafting procedures^[5, 6]^. Allogeneic approaches, including xenogeneic grafts^[7]^, decellularized extracellular matrix scaffolds^[8]^, and synthetic skin substitutes^[8]^, partially alleviate donor shortages; however, these approaches are frequently associated with immune rejection, insufficient vascular integration, poor long-term tissue remodeling, and compromised functional outcomes^[9]^. Therefore, the development of next-generation skin substitutes that simultaneously exhibit high biocompatibility, structural stability, tissue adaptability, and exudate absorption capability has become a central objective in regenerative biomaterials research^[10, 11]^.

Clinically used skin grafts are generally classified into epidermal skin grafts (ESGs), split-thickness skin grafts (STSGs), and full-thickness skin grafts (FTSGs), each imposing distinct structural and functional requirements on biomaterial design. ESGs primarily require rapid epithelial coverage and high surface biocompatibility^[12]^, whereas STSGs demand balanced flexibility, exudate absorption, and tissue integration to support partial dermal regeneration^[13]^. In contrast, FTSGs require substantially enhanced mechanical robustness, structural stability, and long-term remodeling capability to restore the integrity of full-thickness skin tissue^[14]^. Consequently, developing a universal skin substitute capable of simultaneously satisfying these diverse clinical requirements remains highly challenging.

To date, most artificial skin systems have been constructed from extracellular matrix-derived proteins, polysaccharides, or synthetic polymers, including collagen, hyaluronic acid, and poly(lactic-co-glycolic acid) (PLGA). However, these conventional materials often suffer from intrinsic trade-offs among mechanical strength, biodegradability, bioactivity, and fluid absorption capability^[15,16]^. For example, collagen-based materials exhibit excellent cytocompatibility and biological recognition but generally possess poor mechanical stability and rapid degradation behavior due to their weak intermolecular interactions and linear molecular architecture^[17]^. Synthetic polymers can partially improve mechanical performance; however, they frequently compromise biological functionality and tissue adaptability^[18]^. Moreover, extensive chemical crosslinking strategies commonly employed to reinforce these materials may introduce cytotoxicity, structural heterogeneity, and batch-to-batch variability, thereby limiting their translational potential^[18]^.

Protein-based biomaterials, particularly genetically engineered protein systems, have emerged as highly promising candidates for next-generation skin substitutes owing to their intrinsic biocompatibility, programmable molecular architecture, and tunable physicochemical properties^[19, 20]^. Unlike conventional biomaterials, engineered protein matrices enable precise molecular-level regulation of mechanical behavior, degradation kinetics, and biofunctional signaling, thereby providing unique opportunities for the development of structurally adaptive and biologically active artificial skin^[21, 22]^. Nevertheless, the mechanical fragility of protein-based matrices and their strong dependence on chemical crosslinking substantially limit their application in mechanically demanding skin regeneration scenarios.

In recent years, protein-based chain entanglement has emerged as an effective topological strategy for enhancing the mechanical toughness and energy dissipation capability of biomaterials^[23,24]^. Unlike conventional chemically crosslinked networks, entanglement-mediated protein assemblies rely predominantly on physical interactions, thereby reducing dependence on exogenous crosslinking agents while enabling dynamic regulation of intermolecular spatial organization through protein architecture. Such topological interactions can substantially improve mechanical robustness while preserving the intrinsic bioactivity of protein-based matrices. However, excessive entanglement density often leads to highly compact network architectures that restrict chain mobility and reduce matrix adaptability^[25]^. Moreover, protein denaturation and renaturation processes are needed to induce protein-based entanglement for high mechanical materials preparation^[26]^. This structural over-confinement not only compromises network flexibility and stress-relaxation behavior, but may also impair cellular spreading, migration, and matrix remodeling^[27]^. Therefore, although entanglement-mediated protein networks provide a promising route toward mechanically robust biomaterials without extensive chemical crosslinking, precisely balancing network mechanics, structural adaptability, and cytocompatibility remains unresolved in current artificial skin systems^[28]^. To date, few protein-based artificial skin platforms have simultaneously achieved mechanically adaptive reinforcement, cytocompatibility, and spatially integrated regenerative functionality within a single material system.

Here, we establish an entanglement-mediated mechanically adaptive protein network for the construction of bilayer artificial skin with integrated antibacterial and regenerative functionalities. Tandem ferredoxin-like proteins (FLn) were employed as the primary structural scaffold, while flexible molecular linkers were introduced to modulate intermolecular interactions and facilitate stress dissipation throughout the network. Through the synergistic integration of protein chain entanglement, copolymer reinforcement, and photo-triggered crosslinking, a mechanically robust yet structurally adaptive three-dimensional protein matrix was established, enabling precise regulation of network mechanics and matrix flexibility. Building upon this mechanically adaptive platform, a stratified bilayer artificial skin was further engineered through modular biofunctional integration. The epidermal layer was designed to provide antibacterial activity through Zn^2+^-coordinated functional domains, whereas the dermal layer incorporated a collagen-like peptide–epidermal growth factor (CLP–EGF) module to promote cellular proliferation and tissue regeneration. These two functional layers were subsequently integrated through an interfacial in situ assembly strategy, enabling spatial separation of antibacterial and regenerative functions while maintaining coordinated therapeutic activity within a single construct. Collectively, this work establishes a multiscale protein engineering strategy that decouples mechanical reinforcement from structural over-confinement, providing a versatile framework for the development of mechanically resilient and biologically functional protein-based regenerative biomaterials.

## Materials and Methods

### 1. Chemical Reagents and Antibodies

The primary chemical reagents and antibodies used in this study are as follows: All primers and plasmid templates used for gene cloning and plasmid construction were synthesized by Sangon Biotech Co., Ltd. (Shanghai, China). Plasmid construction, amplification, and verification were performed in-house. Molecular biology enzymes, including restriction endonucleases, DNA ligase, and high-fidelity DNA polymerase, were purchased from Thermo Fisher Scientific (USA). Primary and secondary antibodies used in this study are as follows:

Mouse anti-His tag monoclonal antibody (Cat. No. AB18184), β-actin antibody (Cat. No. AB8226), and IL-6 antibody (Cat. No. 1092254-6) were purchased from Abcam (UK). Goat anti-mouse IgG H&L (HRP-labeled, Cat. No. ab205719) was purchased from Abcam (UK). GAPDH antibody (Cat. No. GB15002-100), PI3K antibody (Cat. No. GB11525-100), Phospho-AKT (Ser473) antibody (Cat. No. 150002-100), Total AKT antibody (Cat. No. GB15689-100), IL-10 antibody (Cat. No. GB11108-100), phosphorylated mTOR (Ser2448) antibody (Cat. No. GB114489-100), total mTOR antibody (Cat. No. GB11405-100), and goat anti-rabbit IgG (HRP-labeled, Cat. No. GB23303-100) were purchased from Servicebio Technology (Wuhan, China). The CCK-8 cell viability assay reagent and the live/dead cell dual staining kit were purchased from Beijing Saier Biotechnology Co., Ltd. (China).

### 2. Plasmid Construction, Protein Expression, and Purification

The gene encoding FL8 was codon-optimized and synthesized by Sangon Biotech Co., Ltd. (Shanghai, China). The synthesized fragment was digested with specific restriction enzymes and then directionally cloned into the pET-28a(+) expression vector, which had undergone identical digestion, under the catalysis of T4 DNA ligase^[29]^. The coding sequences for target proteins FL4, FL8-ZINC, and CLP-EGF were assembled and constructed using the overlap extension PCR method. The purified PCR products were digested with specific restriction enzymes and cloned into the pET-28a(+) expression vector, which had undergone identical digestion, using T4 DNA ligase. All recombinant plasmids were validated by DNA sequencing to ensure sequence accuracy.

Recombinant plasmids encoding the target proteins were individually transformed into *Escherichia coli* BL21(DE3) competent cells for heterologous expression^[30]^. Transformed colonies were cultured in Luria–Bertani (LB) medium supplemented with kanamycin (50 μg mL⁻¹) at 37 °C with shaking at 220 rpm until the optical density at 600 nm (OD₆₀₀) reached approximately 0.8. Protein expression was subsequently induced by the addition of isopropyl β-D-1-thiogalactopyranoside (IPTG, final concentration 0.3 mM), followed by incubation at 16 °C for 24–48 h. Cells were harvested by centrifugation and resuspended in lysis buffer. Following sonication-induced cell disruption, the lysates were centrifuged to remove insoluble debris, and the clarified supernatants were collected for protein purification^[31]^. Recombinant proteins carrying an N-terminal 6×His-tag were purified using immobilized metal affinity chromatography (IMAC) with Ni^2+^-charged affinity resin. After extensive washing, bound proteins were eluted using imidazole-containing elution buffer. The purified proteins were subsequently concentrated and buffer-exchanged into deionized water using centrifugal ultrafiltration devices with a molecular weight cutoff of 30 kDa. Protein purity was analyzed by SDS–PAGE followed by Coomassie Brilliant Blue staining. Protein identity was further confirmed by Western blot analysis using an anti-His tag antibody.

### 3. Protein Functionalization and Hydrogel Preparation

#### NHS-PEG-AM-Modified Proteins

To introduce acrylamide (AM) functional groups into proteins, N-hydroxysuccinimide-polyethylene glycol-acrylamide ester (NHS-PEG-AM) ^[32]^was used for covalent modification of the protein’s primary amine groups. The specific procedure is as follows: A solution of NHS-PEG-AM dissolved in dimethyl sulfoxide (DMSO) was mixed with a purified protein solution at a 1:9 volume ratio, ensuring a molar ratio of protein to NHS-PEG-AM of 1:2. The reaction mixture was incubated overnight at 4°C. After reaction completion, transfer the mixture to a pre-chilled ultrafiltration centrifuge tube (30 kDa molecular weight cutoff). Thoroughly remove unreacted NHS-PEG-AM and byproducts by repeated centrifugation and exchange with deionized water. Finally, the purified modified protein solution was freeze-dried to obtain protein-AM lyophilizate, stored at -20°C for later use. Modification success was verified via ¹H NMR spectroscopy.

#### Fabrication of the epidermal layer

Lyophilized FL8-ZINC-AM protein was dissolved in 7 M guanidine hydrochloride to obtain a protein precursor solution at a concentration of 100 mg mL⁻¹. ZnCl₂ was subsequently introduced to a final concentration of 5 mM, followed by the addition of acrylamide monomer to a final concentration of 150 mg mL⁻¹. The precursor solution was incubated at room temperature for 30 min to facilitate homogeneous mixing and preliminary intermolecular assembly. Ammonium persulfate (APS) and the photoinitiator 2-hydroxy-4′-(2-hydroxyethoxy)-2-methylpropiophenone (Irgacure 2959) were then added to final concentrations of 50 mM and 1% (w/v), respectively. The resulting precursor solution was transferred into molds and exposed to 365 nm ultraviolet irradiation for 20 min to induce photopolymerization and network crosslinking, yielding the epidermal-layer hydrogel (FZA).

#### Fabrication of the dermal layer

Lyophilized FL8-AM protein was dissolved in 7 M guanidine hydrochloride at a concentration of 50 mg mL⁻¹. Acrylamide monomer was subsequently added to achieve a final concentration of 75 mg mL⁻¹. Following incubation at room temperature for 30 min, APS and Irgacure 2959 were introduced at final concentrations identical to those used for the epidermal layer (50 mM and 1% (w/v), respectively). The precursor solution was then subjected to 365 nm ultraviolet irradiation for 20 min to complete photopolymerization. The resulting hydrogel was immersed in phosphate-buffered saline (PBS, pH 7.4), and the buffer was refreshed every 2 h over a 24 h period to remove residual guanidine hydrochloride and allow equilibration swelling, yielding the dermal-layer hydrogel (FED).

#### Fabrication of the bilayer artificial skin

The integrated bilayer artificial skin was constructed through a sequential swelling–polymerization strategy. Briefly, the preformed FED hydrogel was equilibrated in PBS prior to assembly. Excess surface liquid was gently removed using filter paper, after which the epidermal precursor solution containing FL8-ZINC-AM, acrylamide, APS, and Irgacure 2959 was uniformly deposited onto the dermal-layer surface. The assembled construct was immediately exposed to 365 nm ultraviolet irradiation for 20 min to induce in situ photopolymerization of the epidermal layer. During this process, interfacial chain interpenetration and network integration occurred simultaneously between the two layers, resulting in a mechanically integrated bilayer artificial skin with distinct structural and functional stratification.

### 4. Physicochemical and Mechanical Characterization of Hydrogel Materials

Prepared monolayer and bilayer artificial skin samples were pre-frozen at -80°C and then freeze-dried in a freeze dryer (LGJ-10C, Foring, China) for 48 h. After gold coating via an ion sputter coater (SC7620, Quorum, UK), dried samples were examined using a field emission scanning electron microscope (SU8010, Hitachi, Japan) at 5 kV acceleration voltage to observe cross-sectional and surface microstructures, with images recorded^[33]^.

Artificial skin samples were cut to appropriate dimensions and adhered to the surface of clean, fresh sheepskin that had undergone depilation. The sheepskin-material composite underwent various deformations including vertical suspension, twisting, flipping, and suturing to qualitatively assess the material’s wet adhesion to biological tissue and suturability. All adhesion and deformation processes were recorded using a high-resolution camera^[34]^.

Mechanical properties of the hydrogels were evaluated using a universal testing machine (Instron 5967, Instron, USA). Hydrogel samples were cut into dumbbell-shaped specimens according to the Chinese National Standard GB/T 1040.3–2006 (Type 3 geometry) and equilibrated in phosphate-buffered saline (PBS) prior to testing. All measurements were performed at room temperature (25 °C) with a constant tensile rate of 5 mm/min. At least five independent samples were analyzed for each group. For cyclic tensile measurements, specimens were subjected to sequential loading–unloading cycles at maximum strains of 25%, 50%, 75%, and 100% to evaluate elastic recovery behavior, hysteresis, and energy dissipation capacity of the network. For uniaxial tensile-to-failure tests, samples were continuously stretched until fracture, and the corresponding stress–strain curves were recorded. Fracture stress, fracture strain, and tensile toughness were subsequently calculated from the obtained mechanical profiles.

After precisely weighing the initial dry weight (W₀) of freeze-dried circular hydrogel samples (approximately 8 mm diameter), immerse them in the following three degradation media^[35]^: (1) phosphate-buffered saline (PBS, pH 7.4), (2) PBS solution containing 50 U mL⁻¹ trypsin, (3) PBS containing 50 U mL⁻¹ of proteinase K. All samples were placed in 1 mL of degradation medium and incubated at 37°C. At predetermined time intervals, 1 μL samples were withdrawn from the mixtures. Protein concentration was measured using a NanoDrop spectrophotometer (Thermo Fisher Scientific) to quantify the accumulated free protein. The protein degradation rate was calculated using the following formula: The cumulative degradation rate (Dₜ) based on protein release at each time point was calculated using the following equation, given the theoretical protein content based on medium volume (V) and initial gel dry weight (W₀): Dt(%) = (Ct × V / W₀ × fp) × 100% where fp is the mass fraction of protein dry weight relative to total monomer dry weight used in hydrogel preparation. Set up at least 3 parallel samples per experimental group.

To investigate biodegradation behavior in vivo, protein components were first labeled with Cy7 NHS ester before hydrogel fabrication. Circular artificial skin constructs (10 mm diameter) containing Cy7-labeled proteins were implanted into full-thickness dorsal skin defects in male BALB/c mice (6–8 weeks old) and secured using interrupted sutures. The degradation process was longitudinally monitored using a small-animal in vivo imaging system at predetermined time points (days 0, 3, 5, 7, and 10 post-implantation). Fluorescence intensity within the implantation region was quantified and normalized to the initial signal at day 0 to evaluate residual material retention and biodegradation kinetics in vivo.

### 5. Evaluation of Antimicrobial Performance

The antibacterial activity of the materials was evaluated using an agar diffusion assay against *E. coli* (ATCC 25922) and *S. aureus* (ATCC 29213). Briefly, 100 μL bacterial suspensions (1 × 10^8^ CFU mL^-1^) were uniformly spread onto LB agar plates. Circular artificial skin samples with identical dimensions were then placed onto the bacterial lawns, while plates without samples served as controls. After incubation at 37 °C for 12 h, inhibition zones surrounding the samples were imaged and quantified by measuring the diameter of the clear zones. Antibacterial activity was evaluated based on the relative inhibition zone size compared with control groups.

To investigate bacterial morphological changes induced by the materials, bacterial samples obtained from both agar diffusion assays and liquid co-culture experiments were collected. Cells were pelleted by centrifugation (8000 rpm, 5 min) and washed twice with PBS (pH 7.4). The bacterial pellets were fixed in 2.5% (v/v) glutaraldehyde for 2 h at room temperature, followed by sequential dehydration in graded ethanol solutions (30%, 50%, 70%, 80%, and 90%, each for 15 min), and three final washes with absolute ethanol. Samples were then subjected to tert-butanol replacement and freeze-dried. The dried specimens were mounted on conductive adhesive, sputter-coated with gold, and imaged using field-emission scanning electron microscopy at an accelerating voltage of 5 kV.

Antibacterial activity under liquid culture conditions was evaluated by monitoring bacterial growth kinetics. Artificial skin samples (0.1 g) were incubated in LB medium containing *E. coli* or *S. aureus* suspensions at an initial concentration of 5 × 10^5^ CFU mL^-1^. Cultures were maintained at 37 °C under shaking conditions (200 rpm). At predetermined time points (0, 2, 4, 6, 12, and 24 h), bacterial growth was quantified by measuring optical density at 600 nm (OD_600_) using a microplate reader. Antibacterial performance was evaluated based on suppression of bacterial growth relative to untreated controls. All experiments were performed in triplicate.

A full-thickness excisional wound (10 mm in diameter) was created on the dorsal skin of anesthetized mice, followed immediately by topical inoculation with 20 μL of bacterial suspension containing either *E. coli* (ATCC 25922) or *S. aureus* (ATCC 29213) at a concentration of approximately 1 × 10^8^ CFU mL^-1^ to establish infected wound models. After bacterial inoculation, wounds were treated with the indicated materials and covered under sterile conditions. At predetermined time points (days 0, 3, and 5 post-infection), wound exudates and surface-associated bacteria were collected using sterile cotton swabs. Each swab was immersed in 1 mL sterile PBS and vortexed thoroughly to detach adherent bacteria. Serial dilutions of the bacterial suspension were prepared, and 100 μL aliquots were spread onto antibiotic-free LB agar plates, followed by incubation at 37 °C for 12 h. CFUs were subsequently counted to quantify bacterial burden at the wound site. Antibacterial efficacy was calculated relative to the untreated control group according to the following equation: Bacteriostatic rate (%) = (1-CFU_experimental_/ CFU_control_) ×100%.

### 6. Cell Compatibility and Cell Behavior Evaluation

The cytocompatibility of the artificial skin constructs was evaluated using a Cell Counting Kit-8 (CCK-8, Beijing Solarbio Science & Technology Co., Ltd.) assay^[36]^. Mouse fibroblasts (L929) and human keratinocytes (HaCaT) were seeded into 96-well plates at a density of 5 × 10³ cells per well and cultured for 24 h to allow cell attachment. The culture medium was subsequently replaced with fresh complete medium containing 3% (w/v) material extracts prepared according to ISO 10993 guidelines. Cells cultured in standard complete medium served as the control group. After 24 h incubation, 10 μL of CCK-8 reagent was added to each well, followed by incubation at 37 °C for 30 min. Absorbance at 450 nm was measured using a microplate reader (Thermo Fisher Scientific). Relative cell viability was calculated using the following equation: Cell viability (%) = (OD_treated_ - OD_blank_) / (OD_control_ - OD_blank_) × 100%. All experiments were performed with at least five independent replicates.

Cell viability on material surfaces was further evaluated using a Calcein-AM/propidium iodide (PI) live/dead staining assay. L929 fibroblasts and HaCaT keratinocytes were seeded into 24-well plates at a density of 2 × 10^4^ cells per well and cultured under the indicated experimental conditions. After treatment, cells were gently rinsed once with PBS to remove residual medium^[37]^. Calcein-AM and PI stock solutions were diluted in PBS at a ratio of 1:2000 according to the manufacturer’s instructions to prepare the staining solution. Samples were incubated with staining solution at 37 °C for 5 min in the dark and subsequently imaged using a fluorescence microscope. Live cells stained green (Calcein-AM), whereas membrane-compromised dead cells stained red (PI).

The effect of material extracts on fibroblast migration was evaluated using an in vitro scratch assay. L929 cells were seeded into 24-well plates at a density of 2 × 10^4^ cells per well and cultured to form confluent monolayers. Linear scratches were generated using a sterile 200 μL pipette tip, followed by three gentle PBS washes to remove detached cells. Cells were subsequently cultured in serum-free medium containing 3% (w/v) extracts of the indicated materials. Serum-free medium containing PBS served as the control. Bright-field images were acquired at identical positions immediately after scratching (0 h) and at 6, 12, and 24 h using an inverted optical microscope. Scratch areas were quantified using Fiji software, and migration efficiency was expressed as wound closure percentage according to the following equation: Healing rate (%) = (1 - A_t_/A₀) × 100%. Where A₀ represents the initial scratch area and A_t_ represents the residual scratch area at the indicated time point. All experiments were performed in triplicate.

Cell–material interactions on dermal-like hydrogels were evaluated using pre-formed dermal-like hydrogels (FED). Hydrogels were placed at the bottom of 6-well plates and equilibrated with complete medium prior to cell seeding. L929 fibroblasts (2 × 10^5^ cells) were resuspended in 20 μL of complete medium and carefully seeded onto the hydrogel surface. After a 2 h incubation to facilitate initial adhesion, 2 mL of pre-warmed complete medium was slowly added along the well walls to avoid detaching attached cells. Cell viability and morphology on the hydrogel were first assessed after 24 h using Calcein-AM/PI staining, following the protocol described in Section 6.2. For long-term proliferation analysis, cells were pre-labeled with the membrane fluorescent dye DiO (5 μM) prior to seeding. Fluorescence images were acquired at fixed fields of view on days 1, 3, and 5 post-seeding using a fluorescence microscope. The number of DiO-positive cells was quantified using ImageJ software, and temporal changes in cell density were used to evaluate proliferative capacity on the hydrogel scaffold.

### 7. Western Blot

Total protein was extracted from L929 cells or mouse wound tissues using a commercial total protein extraction kit (Beyotime, China) according to the manufacturer’s protocol. Protein concentrations were determined using a BCA assay (Beyotime, China). Equal amounts of protein (30 μg per sample) were separated by 12% SDS-PAGE and subsequently transferred onto 0.22 μm polyvinylidene fluoride (PVDF) membranes under wet-transfer conditions. Membranes were blocked with 5% (w/v) non-fat milk in TBST for 1 h at room temperature, followed by incubation with primary antibodies at 4 °C overnight. After extensive washing with TBST, membranes were incubated with horseradish peroxidase (HRP)-conjugated secondary antibodies for 1 h at room temperature. Protein bands were visualized using an enhanced chemiluminescence (ECL) detection system and quantified using ImageJ software. Relative protein expression levels were normalized to GAPDH or β-actin as internal loading controls.

### 8. Hemocompatibility Evaluation

Hemocompatibility was evaluated using a standard hemolysis assay. Fresh whole blood was centrifuged at 1000 rpm to obtain red blood cells (RBCs), which were washed three times with physiological saline and diluted to a 2% (v/v) suspension. Artificial skin samples (5%, w/v) were incubated with 1 mL of RBC suspension at 37 °C for 1 h. Physiological saline and 1% Triton X-100 served as negative and positive controls, respectively. Following incubation, samples were centrifuged at 1000 rpm for 15 min, and hemolysis was assessed by visual inspection of supernatant coloration and by measuring absorbance at 545 nm using a UV–vis spectrophotometer.

### 9. Large-area Skin Trauma Transplantation

All animal experiments were performed in accordance with protocols approved by the Animal Ethics Committee of Inner Mongolia University (SYXK 2020-0006). Male BALB/c mice (4–6 weeks old, 18–22 g; Spefo, Beijing, China) were housed under specific pathogen-free (SPF) conditions with ad libitum access to food and water and acclimatized for 7 days prior to experimentation.

For the establishment of full-thickness skin defect models^[38]^, mice were anesthetized by isoflurane inhalation and the dorsal skin was shaved and sterilized with 75% ethanol. A full-thickness rectangular excision (2 cm × 4 cm), including epidermis, dermis, and part of subcutaneous tissue, was created under sterile conditions. Mice were then randomly assigned to four treatment groups (n ≥ 5 per group): control, commercial dressing (3M^TM^ Tegaderm^TM^), double-layer artificial skin (FBilayer), and cell-laden artificial skin (FBilayer+cell pre-seeded with L929 fibroblasts). Materials were applied to fully cover the wound area and secured with sterile gauze and breathable adhesive film. Dressings were removed on day 3, after which wounds were left exposed for natural healing. Wound healing progression was monitored by photographing the wound bed on days 0, 7, 14, and 21 post-surgery with a scale bar for calibration. Wound areas were quantified using Fiji software, and wound closure rates were calculated as: Wound healing rate (%) = (A_0_ − A_t_) / A_0_ × 100%, where A₀ is the initial wound area and Aₜ is the area at each time point. At day 21, animals were euthanized and wound tissues, together with surrounding normal skin, were harvested for histological analysis. Samples were fixed in 4% paraformaldehyde, embedded in paraffin, and sectioned at 5 μm thickness. Tissue morphology and extracellular matrix remodeling were evaluated using hematoxylin and eosin (H&E) and Masson’s trichrome staining.

### 10. Mouse Diabetic Wound Model Repair

For diabetic wound studies, diabetes was induced in male BALB/c mice via a single intraperitoneal injection of streptozotocin (STZ, 60 mg kg^-1^), freshly prepared in 0.1 M citrate buffer (pH 4.5). Blood glucose levels were measured from tail vein blood using a glucometer beginning 6 days post-injection. Mice with fasting blood glucose levels exceeding 15 mmol L^-1^ for seven consecutive days were considered diabetic and included in subsequent experiments. After stabilization, a 10 mm full-thickness circular excisional wound was created on the dorsal skin of diabetic mice. Animals were randomly divided into four groups (n ≥ 5 per group): diabetic control (saline), ZnO ointment, FBilayer, and FBilayer+cell. Materials were applied immediately after injury to fully cover the wound surface and secured with sterile dressings, which were removed on day 3. Wound closure was documented on days 0, 3, 6, 9, and 12 post-surgery, and wound area was quantified using Fiji software to calculate closure rates using the same formula described above. At day 12, wound tissues were harvested for histological and molecular analyses, including H&E and Masson’s trichrome staining, as well as immunohistochemical and immunofluorescence staining for Ki67, PCNA, PTEN, and related signaling markers.

## Results

### 2.1 Engineering an entanglement-mediated protein network for mechanically adaptive artificial skin

To address the intrinsic conflict between mechanical robustness and structural adaptability in protein-based artificial skin, we engineered an entanglement-mediated protein network through multiscale molecular regulation. Tandem ferredoxin-like proteins containing repetitive folded domains (FL4 and FL8) and dead-mutation of FL8 were designed as the primary structural scaffolds to enable controllable chain entanglement and stress dissipation within the hydrogel matrix **(Fig. 1a)**. Recombinant proteins were successfully expressed and purified, as confirmed by SDS–PAGE and Western blot analysis **(Fig.S1)**. Structural prediction using AlphaFold2 revealed compact folded conformations with repetitive modular architectures **(Fig. 1b)**, providing the structural basis for intermolecular topological interactions. To integrate the engineered proteins into the polymeric network, primary amine groups on the protein surface were functionalized with NHS–PEG–AM, generating photocrosslinkable protein precursors (FL4-PEG-AM and FL8-PEG-AM) **(Fig. 1c)**. Successful modification was confirmed by ¹H NMR spectroscopy **(Fig.S2)**. During subsequent photo-triggered polymerization with acrylamide monomers, adjacent tyrosine residues within the protein chains underwent oxidative coupling to form dityrosine crosslinks, thereby establishing a covalently stabilized entanglement-mediated network **(Fig. 1c)**. To elucidate the essential mechanisms governing hydrogel formation, a series of control experiments were systematically performed **(Fig. 1d)**. Mechanically stable hydrogels were obtained only when three conditions were simultaneously satisfied: (i) incorporation of polymerizable NHS–PEG–AM-modified proteins, (ii) the presence of acrylamide monomers, and (iii) tyrosine-mediated photocrosslinking capability. Mutation of tyrosine residues to alanine completely prevented hydrogel formation, directly confirming that dityrosine coupling serves as the primary crosslinking mechanism underlying network stabilization **(Fig. 1c, d)**. Replacing FL8 with bovine serum albumin (BSA), or removing protein components entirely, abolished stable gel formation, indicating the unique structural contribution of tandem FL proteins to network assembly **(Fig. 1c)**.

**Figure 1.**
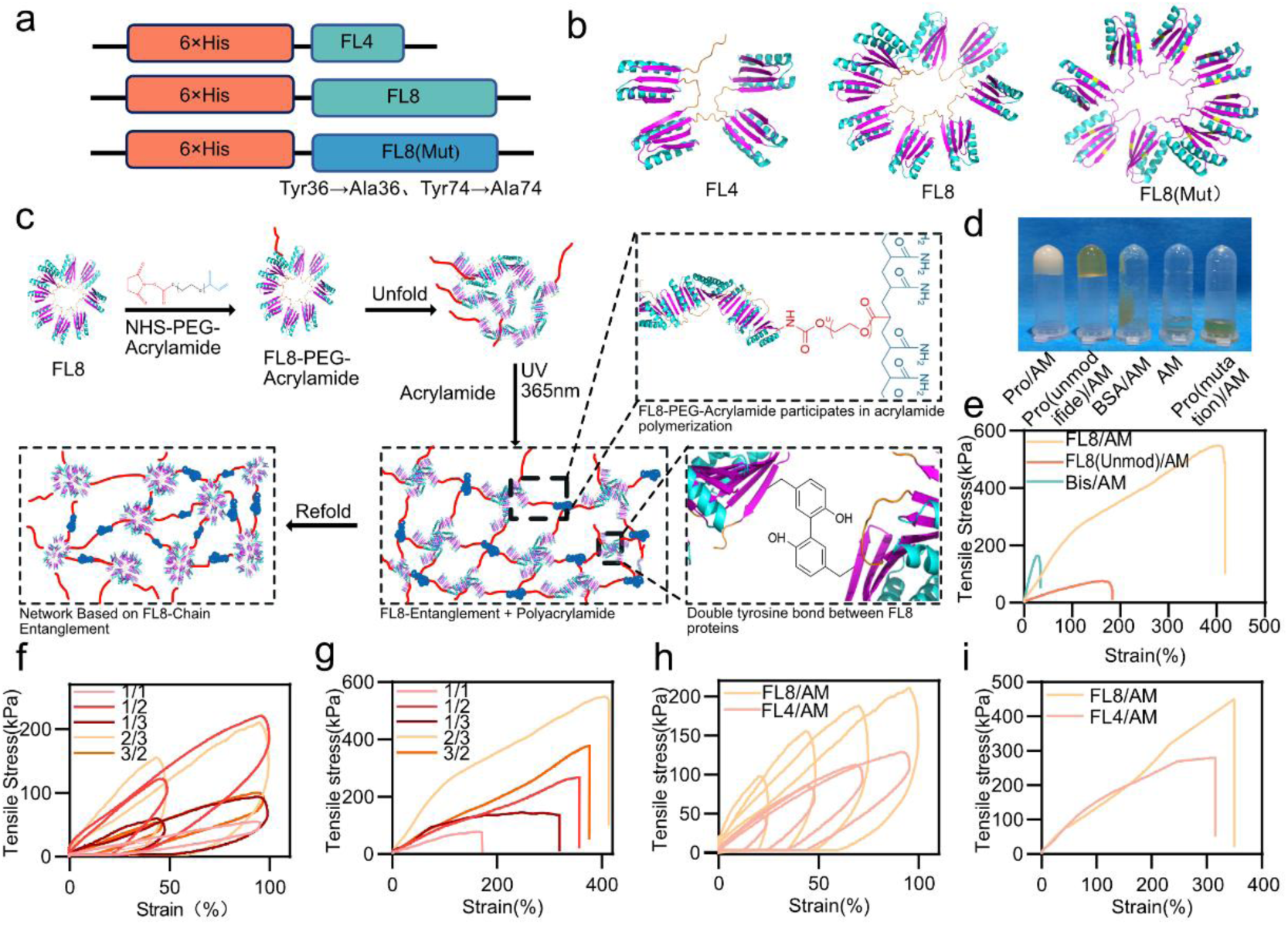
Engineering and mechanistic validation of an entanglement-mediated mechanically adaptive protein network. (a) Schematic illustration of recombinant protein constructs, including tandem ferredoxin-like proteins (FL4 and FL8) and the tyrosine-mutated variant (FL8-Y→A). (b) AlphaFold2-predicted tertiary structures of FL4, FL8, and FL8-Y→A, showing repetitive folded-domain architectures. (c) Schematic illustration of hydrogel network formation. Left, surface functionalization of FL8 proteins using NHS–PEG–AM to generate photocrosslinkable protein precursors. Top right, covalent integration of FL8-PEG-AM into the polyacrylamide network through radical copolymerization. Bottom right, UV-triggered oxidative coupling of adjacent tyrosine residues to form intermolecular dityrosine crosslinks, resulting in stabilization of the entanglement-mediated protein network. (d) Representative photographs of hydrogel formation under different control conditions: complete system FL8-PEG-AM + acrylamide + UV), unmodified FL8 with acrylamide, BSA substituted for FL8, acrylamide alone, and tyrosine-mutated FL8-Y→A. (e) Representative tensile stress–strain curves of hydrogels formed using the complete system and unmodified FL8 proteins without NHS–PEG–AM functionalization. (f, g) Cyclic tensile loading–unloading curves (f) and tensile stress–strain curves to failure (g) of hydrogels prepared with different molar ratios of FL8-PEG-AM to acrylamide. (h, i) Cyclic tensile loading–unloading curves (h) and tensile stress–strain curves to failure (i) of FL4- and FL8-based hydrogels prepared under the optimized formulation condition.

Having established the network formation mechanism, we next investigated the structural parameters governing mechanical adaptation. By systematically varying the ratio between FL8-PEG-AM and acrylamide monomers, we identified an optimal composition that maximized tensile extensibility and toughness **(Fig. 1f, g)**. Hydrogels prepared at a molar ratio of 2:3 exhibited a fracture strain of more than 400% and superior energy dissipation capability **(Fig. 1h, i)**. Furthermore, comparison between FL4- and FL8-based hydrogels revealed that longer protein chains substantially enhanced tensile strength, extensibility, and toughness **(Fig. 1h, i)**. These findings suggest that extended protein architectures promote intermolecular entanglement and dynamic stress dissipation, thereby enabling mechanically robust yet structurally adaptable protein networks suitable for artificial skin applications.

### 2.2 Engineering a bioactive dermal microenvironment through CLP–EGF functional integration

While the entanglement-mediated protein network provided the mechanical foundation for artificial skin construction, effective dermal regeneration additionally requires a bioactive microenvironment capable of supporting cellular proliferation, migration, and matrix remodeling. To endow the mechanically adaptive dermal matrix with regenerative functionality, epidermal growth factor (EGF) was genetically fused with collagen-like peptides (CLP–EGF) to facilitate its stable integration within the protein-based network while preserving extracellular matrix affinity **(Fig. 2a, b)**, Recombinant CLP–EGF was successfully expressed and validated by SDS–PAGE and Western blot analysis **(Fig. S3)**. The purified fusion protein was subsequently incorporated into the entanglement-mediated FL8/polyacrylamide network to construct a bioactive functional epidermal dermal matrix (FED) **(Fig. 2b)**.

**Figure 2.**
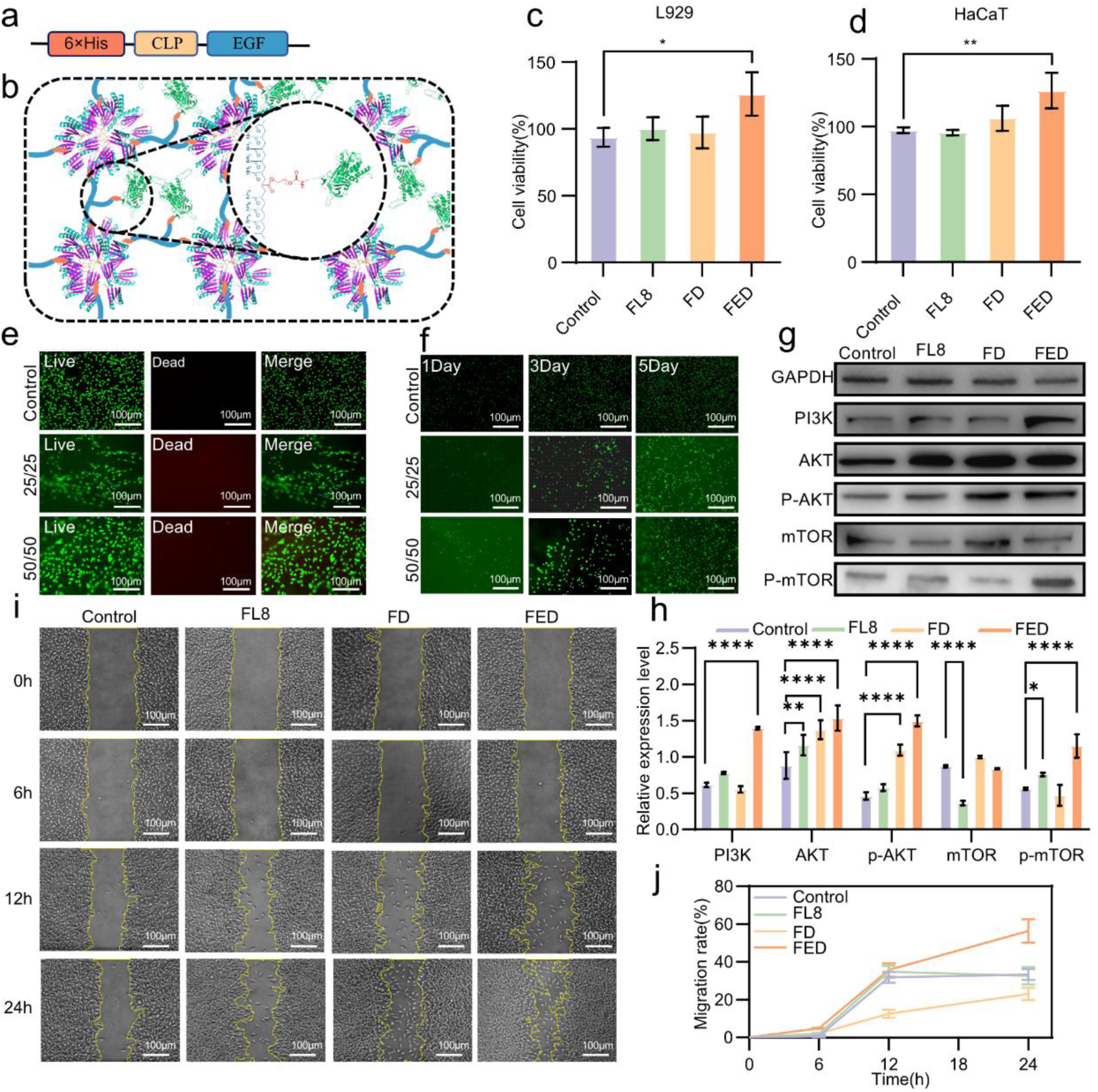
Engineering of a mechanobiologically active dermal matrix through CLP–EGF functional integration. (a) Schematic illustration of CLP–EGF fusion protein design for construction of the bioactive dermal matrix (FED). (b) Schematic illustration of the FED network architecture, showing covalent integration of CLP–EGF within the entanglement-mediated protein matrix. (c, d) CCK-8 analysis of L929 fibroblasts (**c**) and HaCaT keratinocytes (**d**) cultured with FED extracts. (e) Live/dead staining of L929 fibroblasts cultured on FED matrices with different network concentrations. (f) Fluorescence-based proliferation analysis of L929 fibroblasts cultured on FED matrices for 1, 3, and 5 days. (g) Western blot analysis of PI3K–AKT–mTOR signaling in L929 fibroblasts cultured with FED. (h) Quantitative analysis of western blot. (i) Representative migration images at indicated time points. Scale bar, 100 μm. (j) Quantification of wound closure rates. Data are presented as mean ± s.d., n=3, \**P* < 0.05, \*\**P* < 0.01.

We first investigated whether incorporation of CLP–EGF could improve the cytocompatibility and proliferative activity of the mechanically adaptive protein matrix. Extract-based CCK-8 assays demonstrated that FED exhibited excellent cytocompatibility toward both HaCaT keratinocytes and L929 fibroblasts, while significantly enhancing cellular proliferation relative to non-functionalized control matrices **(Fig. 2c, d)**. These findings indicate that incorporation of CLP–EGF effectively transformed the mechanically reinforced protein network into a biologically active regenerative microenvironment.

To further evaluate cell–matrix interactions within the three-dimensional protein network, L929 fibroblasts were directly cultured on FED matrices prepared at different concentrations (25 and 50 mg/mL). Live/dead staining revealed high cellular viability and substantially improved cell attachment on the 50 mg/mL matrix after 24 h of culture **(Fig. 2e)**. Consistently, phalloidin staining showed extensive cytoskeletal spreading and spindle-shaped fibroblast morphology on the higher-concentration matrix **(Fig. S4)**, suggesting that the entanglement-mediated network provided favorable structural support for cell adhesion and cytoskeletal organization. Dynamic proliferation analysis further demonstrated that fibroblasts cultured on the 50 mg/mL FED matrix exhibited significantly enhanced growth over 5 days compared with the lower-concentration group **(Fig. 2f)**. Based on these results, the 50 mg/mL formulation was selected for subsequent biological studies. We next assessed the influence of FED on fibroblast migration, a critical process during dermal remodeling and wound closure. Scratch assays demonstrated that fibroblasts treated with FED extracts exhibited significantly accelerated wound closure rates at 6, 12, and 24 h compared with control groups **(Fig. 2i , j)**. These results indicate that the bioactive dermal matrix not only supports cell viability and proliferation, but also promotes dynamic migratory responses associated with tissue regeneration.

Given the critical role of matrix-regulated signaling in dermal regeneration, we next investigated whether the FED could activate proliferation-associated mechanobiological pathways. Western blot analysis revealed that co-culture with FED markedly increased phosphorylation of AKT and its downstream effector mTOR in L929 fibroblasts relative to controls **(Fig. 2g, h)**, indicating activation of the PI3K–AKT–mTOR signaling axis. As this pathway plays a central role in regulating cellular growth, survival, and metabolic activity, these findings suggest that the CLP–EGF-functionalized matrix actively modulates fibroblast behavior through pro-regenerative intracellular signaling. Collectively, these findings demonstrate that integration of CLP–EGF transforms the entanglement-mediated protein network into a mechanobiologically active dermal microenvironment capable of simultaneously supporting fibroblast adhesion, proliferation, and migration. By coupling mechanically adaptive matrix regulation with regenerative signaling activation, the FED system establishes a functional dermal substitute with substantial potential for full-thickness skin regeneration.

### 2.2 Engineering an antibacterial epidermal barrier through zinc-coordinated protein networks

While the FED layer established a regenerative dermal microenvironment, effective artificial skin systems additionally require an epidermal barrier capable of preventing bacterial invasion during wound healing. To spatially integrate antibacterial functionality within the bilayer architecture, a zinc finger-containing tandem protein variant FL8-ZINC was engineered to construct a zinc-coordinated antibacterial epidermal matrix (FZE) **(Fig. 3a, b)**. Recombinant protein expression was confirmed by SDS–PAGE and His-tag immunoblotting **(Fig. S5)**. Owing to the metal-coordination capability of the zinc finger domain, the engineered protein enabled stable incorporation of Zn^2+^ ions within the entanglement-mediated epidermal network **(Fig. 3b)**. We first evaluated the antibacterial performance of the zinc-functionalized epidermal matrix against both Gram-positive (*S. aureus*) and Gram-negative (*E.coli*) bacteria. Agar diffusion assays demonstrated that Zn^2+^-loaded FZE matrices generated pronounced inhibition zones against both bacterial strains, whereas zinc-free matrices exhibited only weak antibacterial activity and unmodified protein matrices showed negligible inhibition **(Fig. 3c–e)**. To further investigate antibacterial dynamics in a liquid-phase environment, bacterial proliferation kinetics were monitored by measuring optical density (OD_600_) over time. Consistent with solid-phase antibacterial assays, Zn^2+^-functionalized FZE matrices markedly suppressed bacterial growth throughout the incubation period, whereas control groups exhibited rapid proliferation comparable to untreated cultures **(Fig. 3f, g)**. Together, these results demonstrate that incorporation of zinc-coordinated protein domains enables sustained antibacterial regulation within the epidermal layer. Because epidermal antibacterial activity must be balanced with tissue compatibility in artificial skin applications, we next investigated the influence of FZE on mammalian cell viability. CCK-8 assays using HaCaT keratinocytes and L929 fibroblasts revealed moderate suppression of cellular proliferation in Zinc-loaded FZE extracts compared with non-antibacterial control matrices **(Fig. 3h, i)**. Similar trends were observed by live/dead staining analysis **(Fig. S6)**. These findings suggest that although zinc-functionalized epidermal matrices effectively suppress bacterial growth, elevated local Zn^2+^ activity may simultaneously exert concentration-dependent inhibitory effects on mammalian cells. Importantly, such spatially localized antibacterial regulation may be advantageous for bilayer artificial skin systems, as the epidermal layer primarily functions as an external antimicrobial barrier, whereas regenerative cellular activities are predominantly supported by the underlying FED dermal microenvironment. Collectively, these findings demonstrate that zinc-coordinated protein engineering enables construction of an antibacterial epidermal matrix capable of integrating infection-control functionality within the mechanically adaptive bilayer artificial skin platform.

**Figure 3.**
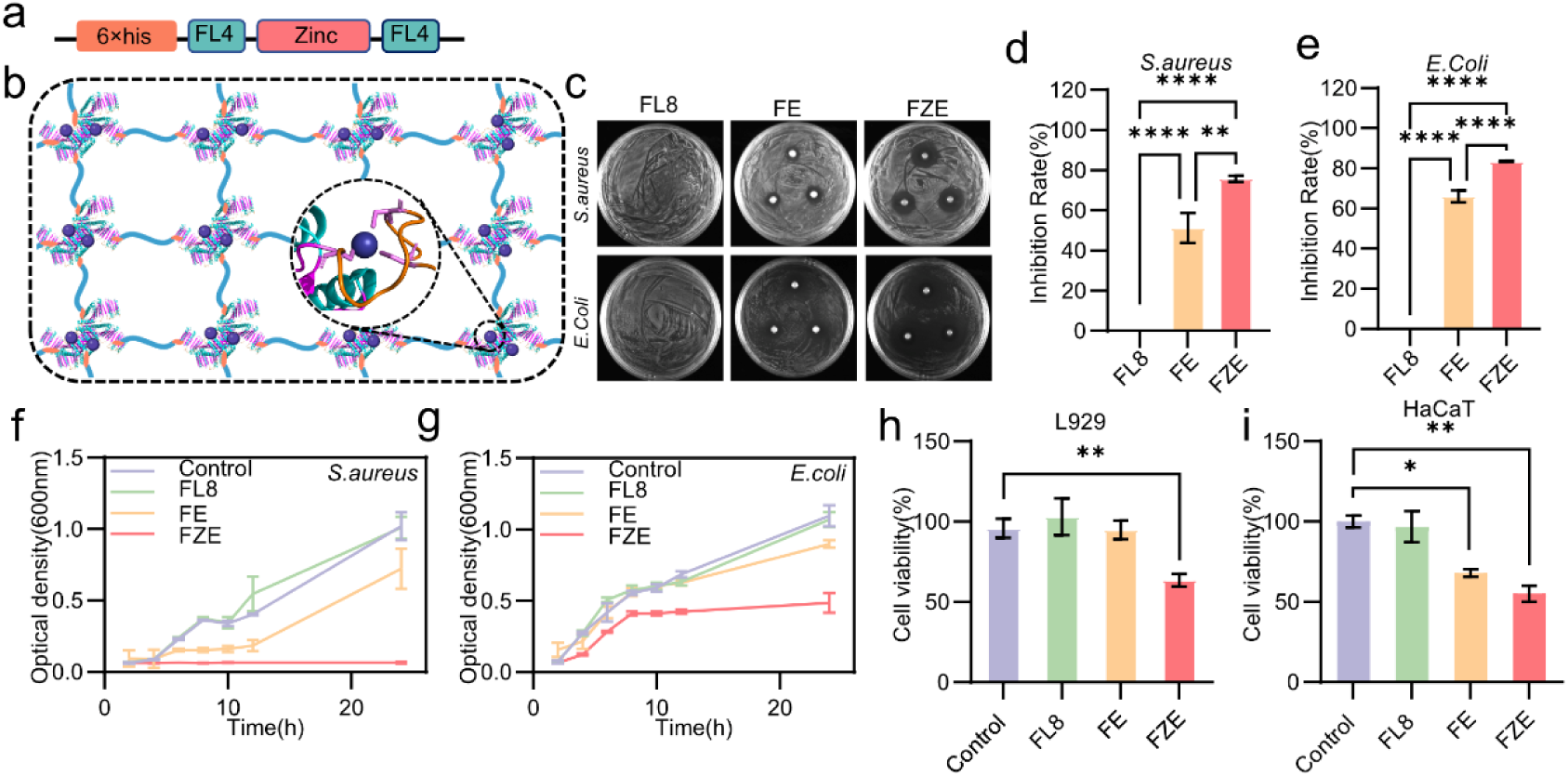
Engineering an antibacterial epidermal barrier through zinc-coordinated protein networks. (a) FL8-ZINC construct containing a zinc finger domain for Zinc coordination within the epidermal matrix. (b) Schematic illustration of the zinc-functionalized epidermal matrix (FZE) and Zn^2+^ coordination within the entanglement-mediated protein network. (c) Representative agar diffusion assays evaluating antibacterial activity. (d, e) Quantification of antibacterial inhibition against *S. aureus* (d) and *E. coli* (e). (f,g) Bacterial growth kinetics of *S. aureus* (f) and *E. coli* (g) cultured with different epidermal matrices, monitored by optical density measurements. (h, i) CCK-8 analysis of L929 fibroblasts (h) and HaCaT keratinocytes (i) cultured with extracts from different epidermal matrices. Data are presented as mean ± s.d., n=3, \**P* < 0.05, \*\**P* < 0.01, \*\*\**P* < 0.001.

### 2.3 Hierarchically compartmentalized bilayer artificial skin enables coordinated mechanical and regenerative functions

Having established a regenerative dermal microenvironment (FED) and an antibacterial epidermal barrier (FZE), we next integrated these two spatially distinct functional compartments into a hierarchically organized bilayer artificial skin through an interfacial in *situ* assembly strategy. In this construct, the FED layer was designed to provide a mechanically compliant and bioactive microenvironment for tissue regeneration, whereas the FZE layer served as a mechanically reinforced antibacterial barrier for protection against external bacterial invasion. The integrated bilayer construct exhibited excellent flexibility, tissue adaptability, and surgical operability. Macroscopically, the artificial skin tolerated repeated twisting, stretching, and bending without structural fracture while maintaining conformal contact with irregular biological tissue surfaces **(Fig. 4a)**. Notably, the construct remained stably attached under inversion and dynamic deformation and could be readily fixed to tissue using standard surgical sutures, indicating favorable applicability for wound coverage and soft-tissue repair. To further investigate the structural organization of the bilayer system, cross-sectional morphologies were characterized by scanning electron microscopy (SEM). Both epidermal and dermal layers displayed interconnected porous architectures with homogeneous network distributions **(Fig. 4b)**, forming a hierarchical structure favorable for nutrient exchange, exudate transport, and cellular infiltration. Importantly, no obvious interfacial delamination or structural discontinuity was observed between the two compartments, suggesting effective interpenetration and interfacial stabilization during in situ polymerization.

**Figure 4.**
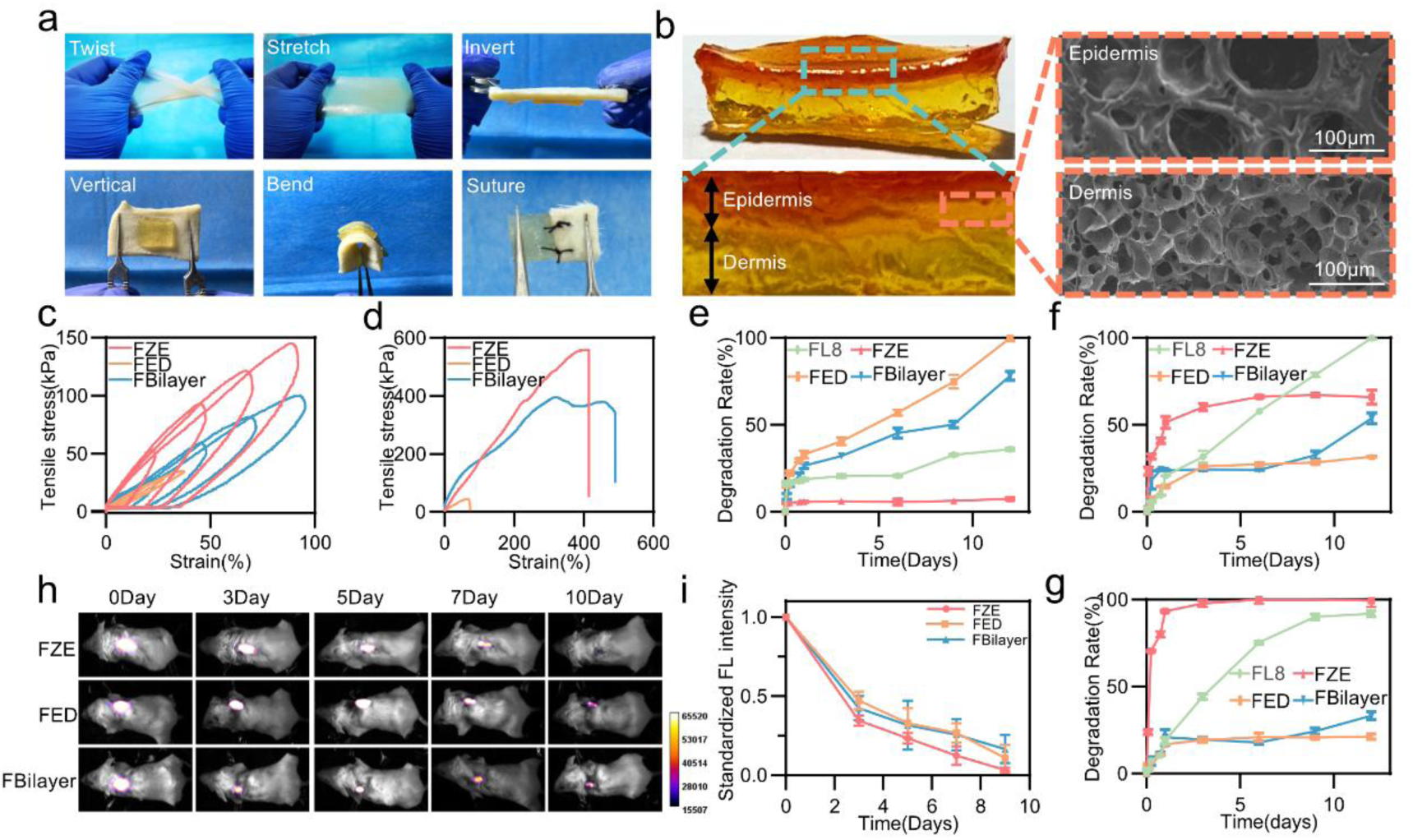
Spatial integration and multiscale regulation of mechanically adaptive bilayer artificial skin. (a) Macroscopic photographs showing the flexibility, deformability, and tissue adaptability of the FBilayer under twisting, stretching, inversion, and suturing conditions. (b) Structural characterization of the bilayer artificial skin. Left, macroscopic cross-sectional image showing the spatially compartmentalized FZE and FED layers. Right, SEM images of the epidermal and dermal compartments showing interconnected porous architectures. Scale bars, 100 μm. (c, d) Mechanical characterization of FED, FZE, and FBilayer. (c) Cyclic tensile loading–unloading curves. (d) Tensile stress–strain curves to failure. (e-g) In vitro degradation profiles of FED, FZE, FBilayer, and unmodified FL8 matrices in PBS (e), proteinase K-containing PBS (f), and trypsin-containing PBS (g). (h, i) In vivo degradation behavior of Cy7-labeled FED, FZE, and FBilayer following subcutaneous implantation in mice. (h) Representative near-infrared fluorescence images at indicated time points. (i) Quantitative analysis of fluorescence intensity over time.

We next examined the independent and cooperative mechanical contributions of the epidermal and dermal compartments. Cyclic tensile loading–unloading and uniaxial tensile tests were performed on FED, FZE, and the integrated bilayer artificial skin (FBilayer) **(Fig. 4c, d)**. The FZE exhibited the highest tensile strength and fracture resistance, likely owing to additional Zn^2+^-mediated coordination reinforcement within the entanglement-mediated protein network. In contrast, the FED dermal matrix displayed a substantially lower elastic modulus, providing a mechanically compliant microenvironment favorable for fibroblast adhesion, proliferation, and matrix remodeling. Because degradation kinetics critically influence the therapeutic performance of regenerative skin substitutes, we next evaluated the biodegradation behavior of the bilayer system under both physiological and enzymatic conditions. In PBS, all matrices exhibited relatively slow degradation profiles **(Fig. 4e–g)**. In contrast, exposure to proteinase K or trypsin markedly accelerated matrix degradation, confirming the enzymatically responsive biodegradability of the protein-based networks. Among all groups, FZE and unmodified FL8 matrices exhibited the most rapid degradation under enzymatic conditions, potentially owing to greater exposure of protein segments within the zinc-functionalized network. By comparison, FED and FBilayer exhibited relatively slower degradation kinetics, suggesting that growth factor integration and multilayer structural organization contributed to enhanced network stabilization during enzymatic hydrolysis. To further evaluate degradation behavior in vivo, Cy7-labeled matrices were subcutaneously implanted in mice and dynamically monitored using near-infrared fluorescence imaging **(Fig. 4h, i)**. All implanted matrices gradually degraded over time, although substantial differences in degradation kinetics were observed among groups. Consistent with in vitro observations, the FZE matrix exhibited the most rapid fluorescence decay, whereas the integrated bilayer artificial skin displayed moderate and temporally sustained degradation behavior. Importantly, substantial signal reduction occurred approximately 10 days post-implantation, corresponding closely to the early proliferative phase of cutaneous wound healing. These findings suggest that the bilayer construct possesses temporally adaptive biodegradation characteristics suitable for transient wound coverage and tissue regeneration.

### 2.4 Spatially compartmentalized bilayer artificial skin enables simultaneous antibacterial protection and regenerative activation

Having established the antibacterial FZE and the regenerative FED, we next investigated whether hierarchical integration of the two functional layers could preserve their respective biological activities within the FBilayer. We hypothesized that spatial compartmentalization within the integrated construct would enable simultaneous antibacterial protection and regenerative activation without mutual functional interference. To evaluate whether the antibacterial function of the epidermal compartment was retained after bilayer integration, antibacterial activity against *E.coli* and *S.aureus* was first assessed using agar diffusion assays. Both FZE and FBilayer generated distinct inhibition zones against Gram-negative and Gram-positive bacteria, whereas negligible antibacterial activity was observed in the FED group **(Fig. 5a–c)**. Importantly, the antibacterial efficacy of FBilayer remained comparable to that of the standalone FZE matrix, indicating that incorporation of the regenerative dermal compartment did not compromise epidermal antibacterial performance. To further examine bacterial responses following exposure to the antibacterial matrices, bacterial morphologies near the inhibition boundary were analyzed by SEM. Bacteria treated with FZE or FBilayer exhibited pronounced membrane collapse, surface wrinkling, and structural deformation, whereas untreated bacteria maintained intact and smooth morphologies **(Fig. 5a)**. Consistent with solid-phase antibacterial assays, bacterial growth kinetics in liquid culture demonstrated that FBilayer significantly suppressed the proliferation of both E. coli and S. aureus throughout the monitoring period, exhibiting antibacterial performance comparable to FZE alone **(Fig. 5d, e)**.

**Figure 5.**
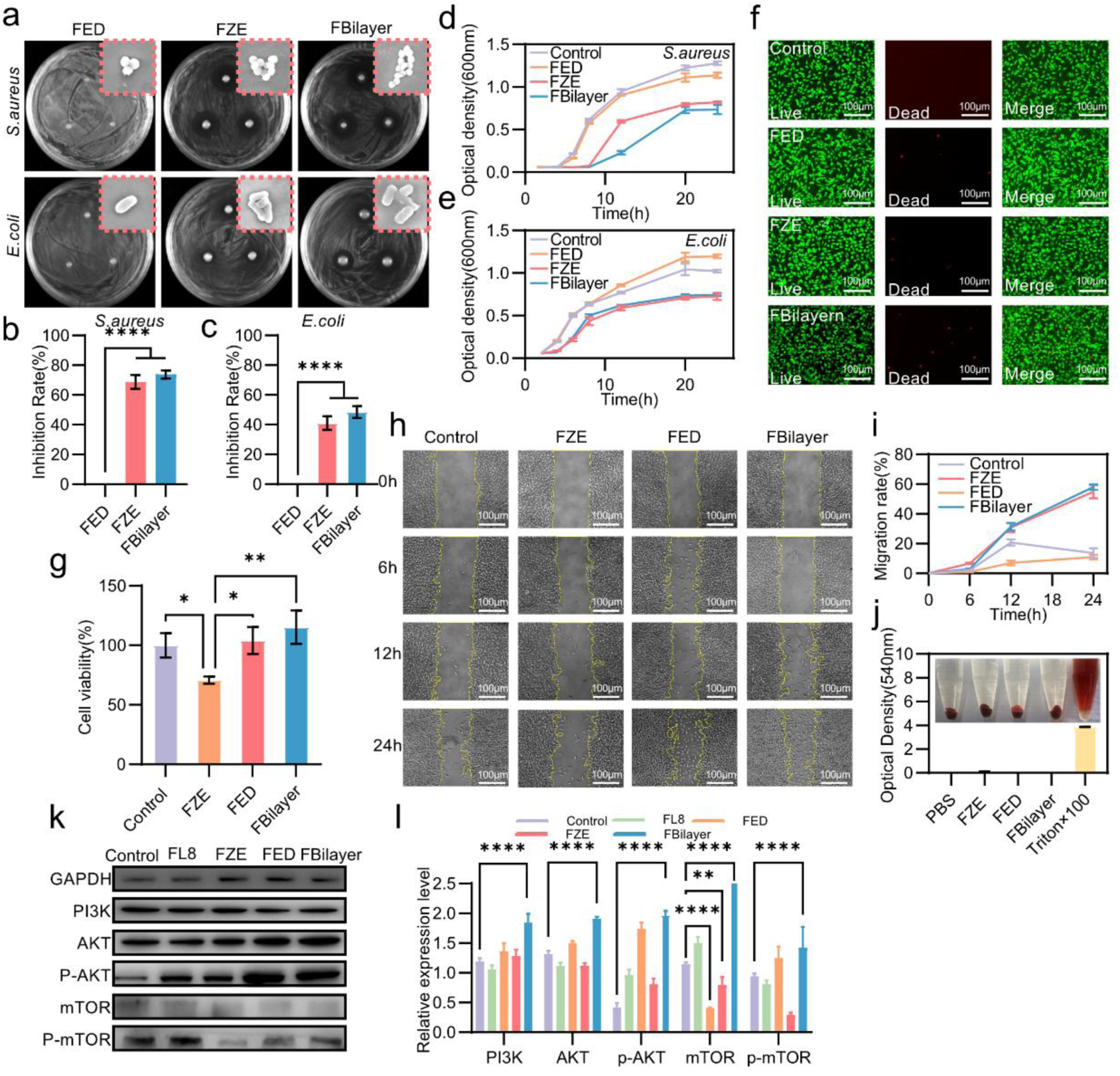
Spatially compartmentalized bilayer artificial skin enables simultaneous antibacterial protection and regenerative activity. (a) Agar diffusion assays evaluating antibacterial activity of FED, FZE, and FBilayer against *S.aureus* and *E.coli*, together with SEM images showing bacterial morphologies following treatment with different matrices. Scale bar, 200 μm. (b, c) Quantification of antibacterial inhibition against *S. aureus* (b) and *E. coli* (c). (d, e) Bacterial growth kinetics of *S. aureus* (d) and *E. coli* (e) cultured with different matrices, monitored by optical density measurements (OD_600_). (f) Live/dead staining of L929 fibroblasts cultured with different matrices for 24 h. Scale bar, 100 μm. (g) CCK-8 analysis of L929 cell viability following co-culture with extracts from different matrices. (h,i) Scratch wound assays evaluating L929 cell migration following treatment with different matrices. (h) Representative scratch images at indicated time points. Scale bar, 100 μm. (i) Quantification of wound closure rates. (j) Hemolysis assay evaluating blood compatibility of FED, FZE, and FBilayer following incubation with mouse red blood cells. Representative photographs of supernatants after centrifugation are shown. (k,l) Western blot analysis of PI3K–AKT–mTOR signaling in L929 cells treated with different matrices. (k) Representative immunoblots. (l) Quantification of p-AKT/AKT and p-mTOR/mTOR expression levels normalized to GAPDH. Data are presented as mean ± s.d. \**P* < 0.05, \*\**P* < 0.01, \*\*\**P* < 0.001, \*\*\*\**P* < 0.0001; n.s., not significant.

We next investigated whether integration of the antibacterial epidermal layer affected the cytocompatibility and regenerative activity of the dermal compartment. Live/dead staining and CCK-8 assays demonstrated that FBilayer maintained favorable compatibility toward both L929 fibroblasts and HaCaT keratinocytes **(Fig. 5f, g and Fig. S7)**. Compared with the highly bioactive FED matrix, FBilayer exhibited slightly reduced proliferative activity, likely owing to the localized antibacterial influence of the zinc-functionalized epidermal layer. Nevertheless, the bilayer constructs still supported robust cell survival and proliferation, indicating that hierarchical compartmentalization enabled effective balancing between antibacterial activity and cytocompatibility. We further evaluated the effect of FBilayer on cellular migration behavior using scratch assays. Both FBilayer and FED significantly accelerated wound closure compared with control and FZE groups **(Fig. 5h, i)**, indicating that the regenerative functionality of the dermal compartment was largely preserved following bilayer integration. To assess blood compatibility, hemolysis assays were performed using red blood cells co-incubated with different matrices. FED, FZE, and FBilayer all exhibited minimal hemolytic activity comparable to the PBS negative control, confirming favorable hemocompatibility of the protein-based matrices **(Fig. 5j)**. We next investigated whether the regenerative signaling activity observed in FED could be maintained after hierarchical integration into the bilayer construct. Western blot analysis demonstrated that treatment with FBilayer significantly increased phosphorylation of AKT and mTOR in L929 cells, consistent with activation of the PI3K–AKT–mTOR signaling pathway observed in the FED group **(Fig. 5k, l)**. These findings suggest that the regenerative signaling capability of the dermal compartment remained functionally preserved within the integrated bilayer architecture. Collectively, these results demonstrate that hierarchical spatial integration enables the bilayer artificial skin to simultaneously maintain antibacterial protection, regenerative bioactivity, cytocompatibility, and hemocompatibility within a unified construct. More importantly, the compartmentalized bilayer architecture effectively balances the intrinsic conflict between antibacterial activity and tissue regenerative capacity, providing a promising design framework for multifunctional regenerative skin substitutes.

### 2.5 Hierarchically integrated bilayer artificial skin enables infection-resistant large-area skin replacement and regenerative repair in vivo

We next investigated whether the FBilayer could coordinately regulate infection control, tissue regeneration, and large-area skin replacement in vivo using mice wound models. To first evaluate antibacterial performance under physiologically relevant conditions, infected full-thickness wound models were established using *S.aureus* and *E.coli*. Circular full-thickness skin defects (1 cm diameter) were generated on the dorsal region of mice, followed by local bacterial inoculation and treatment with saline, ZnO nanoparticles, commercial wound dressing (3M Tegaderm^TM^), FBilayer, or fibroblast-seeded FBilayer (FBilayer+cell). Wound bacterial burden was quantified by colony-forming unit (CFU) analysis at postoperative days 0, 3, and 5 **(Fig. 6a–d)**. Compared with saline and commercial dressing groups, FBilayer-treated wounds exhibited substantially reduced bacterial colonization throughout the observation period. Importantly, the antibacterial efficacy of FBilayer remained comparable to that of cell-loaded FBilayer, indicating that the intrinsic antibacterial functionality of the epidermal compartment was sufficient to maintain infection control in vivo. By comparison, ZnO-treated wounds exhibited relatively limited antibacterial efficacy under the same experimental conditions. These findings demonstrate that the zinc-functionalized epidermal barrier effectively suppresses bacterial proliferation within complex wound microenvironments, thereby preserving a more favorable regenerative niche for subsequent tissue repair.

**Figure 6.**
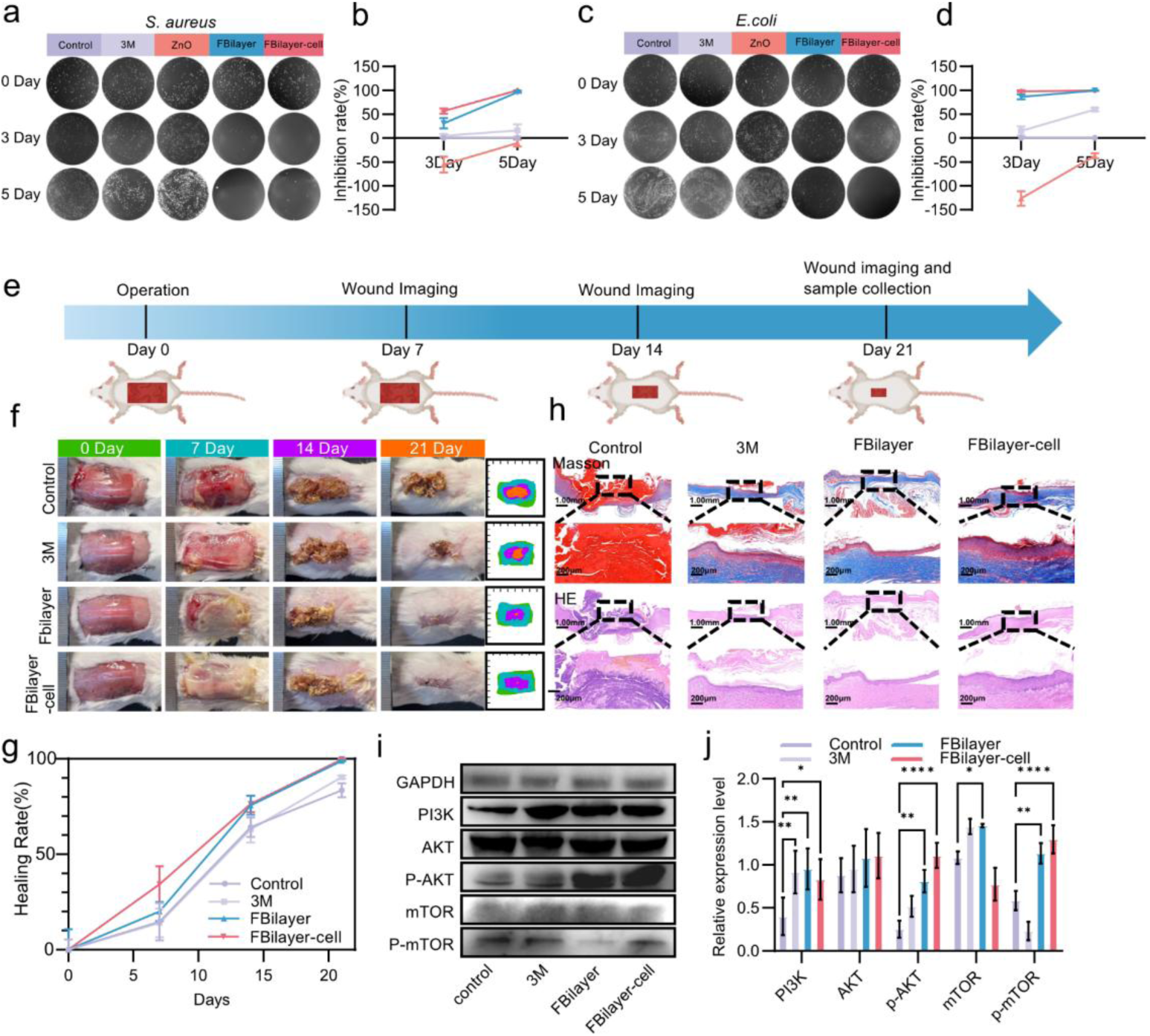
In vivo antibacterial efficacy and transplantation-scale skin regeneration of the bilayer artificial skin. (a) Representative agar plates of *S. aureus* colonies collected from wound exudates at days 0, 3, and 5 post-treatments. (b) Quantification of colony-forming units (CFUs). (c) Antibacterial activity in an *Escherichia coli*-infected wound model. Representative bacterial colony plates. (d) Quantification of CFUs at indicated time points. (e) Schematic illustration of the extensive full-thickness skin defect model (2 × 4 cm) and treatment procedure. (f) Representative photographs of wound repair following transplantation-scale skin replacement at days 0, 7, 14, and 21. Scale bar, 1 cm. (g) Quantification of wound closure rates over time. (h) Histological evaluation of regenerated tissue at day 21 by H&E and Masson’s trichrome staining. Scale bars, 200 μm. (i) Western blot analysis of PI3K–AKT–mTOR pathway activation in regenerated wound tissue at day 21. (j) Quantification of protein expression levels normalized to GAPDH. Data are presented as mean ± s.d. n = 3, \**P* < 0.05, \*\**P* < 0.01.

We next investigated whether FBilayer could function as a transplantable artificial skin substitute for large-area tissue replacement in vivo. To establish a clinically challenging regenerative model, an extensive full-thickness dorsal skin defect (2 cm × 4 cm; approximately 8 cm^2^) was created in mice, corresponding to nearly 40% of the total dorsal skin area **(Fig. 6e)**. Such large-area defects exhibit severely impaired spontaneous healing capacity and therefore represent a stringent model for evaluating functional skin substitutes under high regenerative burden. Animals were treated with saline, commercial wound dressing, FBilayer, or fibroblast-seeded FBilayer (FBilayer+cell), and wound repair was dynamically monitored over 21 days using serial macroscopic imaging **(Fig. 6f, g)**. Notably, the implanted FBilayer remained stably integrated within the wound bed throughout the transplantation period and maintained continuous coverage of the defect area during tissue regeneration. Compared with saline and commercial dressing groups, FBilayer-treated wounds exhibited significantly accelerated wound closure and tissue reconstruction, with near-complete epithelialization achieved by day 21 **(Fig.6 f, g)**. Importantly, the implanted bilayer construct survived and maintained structural integrity throughout the regenerative process, demonstrating its capability to function as a mechanically stable and biologically active artificial skin substitute for large-area tissue replacement. Pre-seeding with fibroblasts did not produce substantial additional improvement in wound closure kinetics, suggesting that the intrinsic regenerative activity of the bilayer construct itself plays a dominant role during tissue repair. To further evaluate the quality of regenerated tissue, histological analyses were performed on wound sections collected at day 21 post-injury **(Fig. 6h)**. Hematoxylin and eosin (H&E) staining demonstrated that FBilayer-treated wounds developed continuous epidermal structures with more organized dermal architectures compared with control groups. In addition, regenerated tissues in the FBilayer group exhibited improved epidermal continuity, enhanced dermal thickness, and more mature extracellular matrix organization, indicating superior structural reconstruction following transplantation **(Fig. 6h)**. Masson’s trichrome staining further revealed substantially enhanced collagen deposition and collagen fiber alignment within regenerated tissues after FBilayer treatment, suggesting accelerated extracellular matrix remodeling and improved tissue maturation **(Fig. 6h)**. To investigate whether the regenerative signaling activity observed in vitro was preserved during in vivo wound healing, we next analyzed activation of the PI3K–AKT–mTOR pathway in wound-edge tissues. Western blot analysis demonstrated that FBilayer treatment significantly increased phosphorylation of AKT and mTOR compared with saline and commercial dressing groups **(Fig. 6i, j)**, consistent with enhanced proliferative and regenerative signaling during tissue repair. These findings suggest that the regenerative dermal compartment remains biologically active following implantation. We additionally examined expression of IL-6 and IL-10 within wound-edge tissues at day 7 post-surgery **(Fig. S8)**. No substantial differences in IL-6 or IL-10 expression were observed among FBilayer, saline, and commercial dressing groups, suggesting that the accelerated regenerative effects of FBilayer are not primarily associated with broad modulation of these canonical inflammatory cytokines under the present experimental conditions. Collectively, these findings demonstrate that the hierarchically integrated bilayer artificial skin enables infection-resistant large-area skin replacement and regenerative repair in vivo, highlighting its strong potential as a mechanically adaptive and biologically active artificial skin substitute for extensive skin transplantation and complex wound reconstruction.

### 2.5 FBilayer reverses regenerative impairment in diabetic wounds through sustained activation of pro-regenerative signaling

To further investigate the therapeutic potential of FBilayer under pathological healing conditions, we next evaluated its regenerative performance in a streptozotocin (STZ)-induced diabetic wound model characterized by impaired tissue repair and chronic regenerative deficiency **(Fig. 7a)**. Stable hyperglycemia was established through repeated STZ administration, resulting in sustained blood glucose levels of approximately 15 mmol/L **(Fig. S9)** prior to wound creation. Full-thickness dorsal skin defects (1 cm diameter) were subsequently generated in diabetic mice, followed by treatment with saline, ZnO ointment, FBilayer, or fibroblast-seeded FBilayer (FBilayer+cell). Wound healing progression was dynamically monitored over 12 days using serial macroscopic imaging and quantitative wound closure analysis (Fig. 7b, c). Compared with saline and ZnO-treated groups, FBilayer-treated wounds exhibited markedly accelerated healing kinetics throughout the observation period. By day 12, wounds treated with FBilayer had undergone near-complete epithelialization and displayed substantially improved tissue closure. Similar therapeutic outcomes were observed in the FBilayer+cell group, further suggesting that the intrinsic bioactivity of the bilayer construct itself is sufficient to support diabetic wound regeneration.

**Figure 7.**
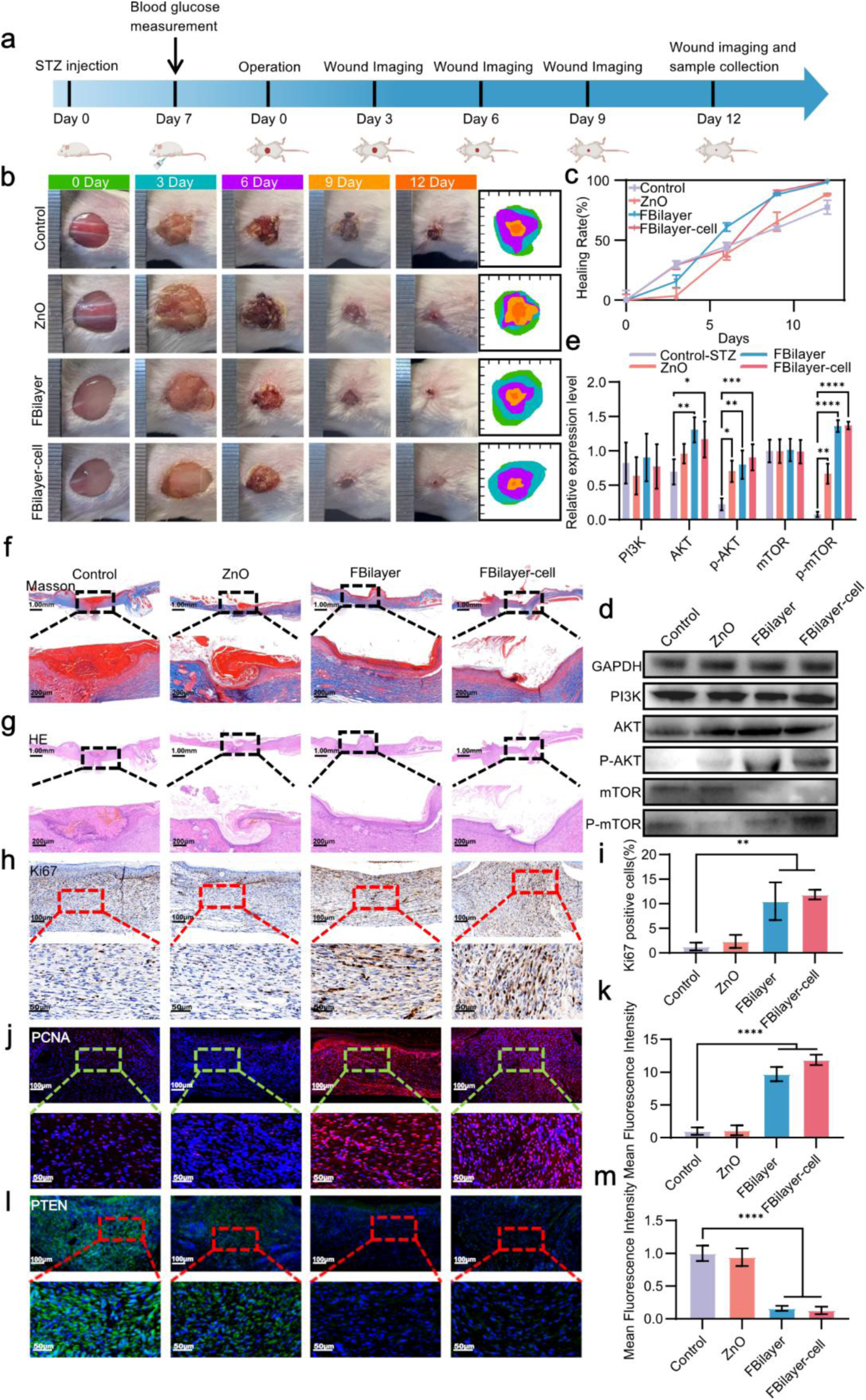
FBilayer accelerates diabetic wound repair through activation of regenerative signaling pathways. (a) Schematic illustration of the diabetic wound-healing model and treatment workflow. Diabetes was induced by streptozotocin (STZ) administration, followed by creation of full-thickness dorsal skin defects and treatment with saline, ZnO ointment, FBilayer, or FBilayer+cell constructs. (b) Representative macroscopic images of wound closure during healing. (c) Quantification of wound closure rates over 12 days. FBilayer-treated wounds exhibited significantly accelerated closure compared with control and ZnO groups. Data are presented as mean ± s.d., n = 3. \**P* < 0.05, \*\**P* < 0.01. (d, e) Western blot analysis and quantification of PI3K–AKT–mTOR pathway activation in wound tissues at day 12. FBilayer treatment markedly increased phosphorylation of AKT and mTOR. GAPDH served as the loading control. Data are presented as mean ± s.d., n = 3, \**P* < 0.05, \*\**P* < 0.01. (f, g) Masson’s trichrome and H&E staining of regenerated tissues at day 12. Scale bars, 1 mm. (h, i) Ki67 immunohistochemical staining and quantification of proliferating cells within regenerated tissues. FBilayer significantly increased Ki67-positive cell populations. Scale bars, 100 μm. Data are presented as mean ± s.d. n = 3, \*\**P* < 0.01. (j, k) Representative PCNA immunofluorescence images and fluorescence quantification of wound tissues. FBilayer treatment enhanced proliferative activity during diabetic wound repair. Scale bars, 100 μm. Data are presented as mean ± s.d. n = 3, \*\**P* < 0.01. (l, m) PTEN immunofluorescence staining and quantitative analysis. FBilayer-treated wounds exhibited reduced PTEN expression relative to control groups, consistent with activation of PI3K–AKT signaling. Scale bars, 100 μm. Data are presented as mean ± s.d. n = 3, \*\**P* < 0.01.

To evaluate the quality of regenerated tissues, histological analyses were performed on wound sections harvested at day 12 post-injury **(Fig. 7f–k)**. H&E staining demonstrated that FBilayer treatment resulted in thinner and more continuous neo-epidermal structures together with more mature dermal organization compared with control groups. Masson’s trichrome staining further revealed enhanced collagen deposition and improved extracellular matrix organization within regenerated tissues, indicating accelerated tissue remodeling and structural restoration under diabetic conditions.

Because impaired cellular proliferation represents a central pathological feature of diabetic wound healing, we next assessed proliferative activity within regenerated tissues. Ki67 immunohistochemical staining and PCNA immunofluorescence analysis demonstrated significantly increased proliferative cell populations in FBilayer-treated wounds compared with saline and ZnO groups **(Fig. 7h–k)**, indicating that the bilayer artificial skin effectively restores regenerative cellular activity within the diabetic wound microenvironment.

To further elucidate the molecular mechanisms underlying this regenerative rescue effect, we analyzed activation of the PI3K–AKT–mTOR signaling pathway in wound tissues **(Fig. 7d, e)**. FBilayer treatment significantly increased phosphorylation levels of AKT and mTOR relative to control groups, consistent with activation of pro-survival and pro-proliferative signaling during tissue regeneration. In parallel, immunofluorescence analysis demonstrated significantly reduced expression of PTEN, a major negative regulator of PI3K–AKT signaling, in FBilayer-treated wounds **(Fig. 7l, m)**. These findings collectively indicate that FBilayer restores regenerative capacity in diabetic wounds through sustained activation of the PI3K–AKT–mTOR pathway.

Chronic inflammation is closely associated with diabetic wound pathology, we additionally evaluated expression of IL-6, IL-10, macrophage polarization markers (CD86/CD206), and TGF-β within regenerated tissues **(Fig. S10 and S11)**. No substantial differences were observed among treatment groups, suggesting that the therapeutic effects of FBilayer are not primarily mediated through broad immunomodulatory remodeling under the present experimental conditions. Instead, the regenerative activity of FBilayer appears to predominantly arise from direct enhancement of cellular proliferation and survival signaling together with infection-resistant wound stabilization. Collectively, these findings demonstrate that FBilayer effectively reverses regenerative impairment in diabetic wounds by restoring proliferative signaling and tissue regenerative activity under pathological conditions. These results further highlight the strong translational potential of this hierarchically engineered artificial skin platform for the treatment of chronic and difficult-to-heal wounds.

## Discussion

The development of artificial skin substitutes capable of simultaneously achieving mechanical robustness, biological functionality, and transplantation adaptability remains a longstanding challenge in regenerative biomaterials^[39]^. Here, we establish a topology-mediated protein engineering strategy for the construction of hierarchically integrated bilayer artificial skin, enabling coordinated regulation of mechanics, antibacterial protection, and regenerative activation within a single biosynthetic protein platform. By integrating entanglement-governed network mechanics with spatially compartmentalized biofunctionalization, this system supports large-area skin replacement and regenerative repair under both acute and pathological wound conditions.

A central conceptual advance of this work lies in the introduction of protein chain entanglement as a mechanically adaptive topological framework for regenerative biomaterials^[25]^. Conventional protein-based hydrogels frequently rely on dense chemical crosslinking to improve structural stability; however, excessive covalent fixation often compromises matrix flexibility, stress relaxation, and cellular adaptability^[40]^. In contrast, the present system utilizes long-chain engineered protein assemblies to establish entanglement-mediated physical interactions that cooperate with photo-triggered intermolecular crosslinking, thereby generating a mechanically resilient yet dynamically deformable protein network. Importantly, this topology-mediated architecture enables efficient energy dissipation under mechanical loading while preserving matrix adaptability required for cellular remodeling and tissue integration^[41]^.

This work further demonstrates the importance of spatially compartmentalized functional integration in artificial skin engineering. Native skin exhibits highly stratified structural and biological organization, in which epidermal and dermal compartments perform distinct yet coordinated physiological functions^[42]^. Inspired by this hierarchical architecture, we engineered a zinc-functionalized antibacterial epidermal layer together with a CLP–EGF-integrated regenerative dermal layer, thereby enabling functional decoupling of antimicrobial protection and regenerative activation within a unified construct. Unlike conventional homogeneous wound dressings^[43]^, this bilayer strategy more closely recapitulates the compartmentalized functional organization of native skin tissue while preserving cooperative interlayer interactions during repair.

Notably, previous protein-entangled biomaterials have primarily focused on mechanically static tissue substitutes, such as cartilage-like constructs, and often require extensive denaturation–renaturation processes that compromise protein bioactivity^[24, 44]^. In the present study, incorporation of flexible PEG-mediated molecular linkers further alleviated interfacial rigidity between protein and polymer phases, promoting more homogeneous stress transfer throughout the network. This strategy transformed conventional physically entangled assemblies into a physically–chemically cooperative interpenetrating system with improved mechanical coordination and enhanced preservation of protein functionality. Such multiscale regulation partially overcomes the long-standing trade-off between mechanical strength and biological activity that commonly limits protein biomaterials.

Beyond mechanical regulation, this work further highlights the importance of spatially compartmentalized functional integration in artificial skin engineering. Native skin exhibits intrinsically stratified structural and biological organization, in which epidermal and dermal compartments perform distinct yet cooperative physiological functions. Inspired by this hierarchical architecture, we engineered a zinc-functionalized antibacterial epidermal layer together with a CLP–EGF-integrated regenerative dermal layer, enabling functional decoupling of antimicrobial protection and regenerative activation within a unified construct. Compared with homogeneous hydrogel dressings, this bilayer strategy more closely recapitulates the compartmentalized organization of native skin while preserving cooperative interfacial integration during tissue repair.

Importantly, the FBilayer construct demonstrated transplantation adaptability for extensive skin replacement under high regenerative burden. The material successfully supported stable coverage and survival within full-thickness skin defects approaching ∼40% of the dorsal surface area in mice (∼8 cm^2^), indicating substantial structural stability during large-area tissue remodeling. Unlike conventional hydrogel dressings primarily designed for temporary wound coverage, the present system maintained mechanical integrity, tissue adhesion, and regenerative support throughout the repair process. Moreover, the protein-engineered construct gradually degraded in synchrony with tissue regeneration, thereby avoiding persistent foreign-body coverage commonly associated with nondegradable polymer dressings. These findings suggest that entanglement-mediated protein engineering may provide a promising route toward transplantation-capable artificial skin substitutes for extensive tissue reconstruction.

The regenerative potential of this platform was further validated in diabetic wound models characterized by impaired proliferation and chronic healing dysfunction^[45]^. FBilayer treatment significantly accelerated wound closure, enhanced extracellular matrix remodeling, and activated PI3K–AKT–mTOR signaling within diabetic wound tissues. Interestingly, minimal alterations were observed in classical inflammatory cytokines or macrophage polarization markers, suggesting that the primary therapeutic mechanism of FBilayer is not dependent on broad immunoregulatory remodeling. Instead, the material appears to establish an infection-resistant regenerative microenvironment that directly supports proliferative and pro-survival signaling activation. This mechanism differs from many immunomodulatory biomaterials^[46]^ and may provide complementary therapeutic opportunities for chronic wound repair.

Despite these promising findings, several limitations remain. First, although the present study demonstrates successful skin replacement and regenerative repair in murine models, long-term evaluation in large-animal systems will be necessary to further assess vascular integration, scar remodeling, and mechanical durability under clinically relevant conditions. Second, while the current platform integrates antibacterial and regenerative functionalities, future optimization may enable incorporation of additional bioactive modules, including angiogenic, neurodegenerative, or immunoregulatory components. Finally, the intrinsic modularity and genetic programmability of engineered protein systems may further facilitate scalable biosynthesis and personalized functional customization for diverse regenerative applications.

In summary, this work establishes a topology-mediated protein engineering framework for the development of mechanically adaptive and biologically functional artificial skin substitutes. By integrating entanglement-governed mechanics, compartmentalized biofunctionality, and regenerative signaling activation within a hierarchically organized protein network, this strategy provides a promising platform for large-area skin transplantation and complex wound repair. More broadly, these findings highlight the emerging potential of programmable protein biomaterials as next-generation regenerative systems for translational tissue engineering and bioactive artificial organs.

## Supporting Information

The Supporting Information is available free of charge at https://

## Acknowledgements

The animal experiments were approved by Inner Mongolia University (SYXK 2020-0006). This project was supported by a grant from Inner Mongolia Autonomous Region Science and Technology Project (2023YFHH0010); National Natural Science Foundation (22365022, 82304409); Inner Mongolia Natural Science Foundation (2026SHZR3474); All authors have approved the final version of this manuscript. We wish to thank the Electron Microscopy Centre of Inner Mongolia University for the microscopy and microanalysis of our specimens.

## Author Contributions

CRediT: Liping Wang and Yinan Sun designed the study and drafted the manuscript; Xing Liu was responsible for the conception of the study; Ruoxuan Wang was responsible for data acquisition; Jinxia Huang was responsible for data interpretation; Wenbo Wang was responsible for validation; Kongxi Fan was responsible for visualization; Jia Bai was responsible for visualization; Zhiying Dong was responsible for validation; Shuang Jia was responsible for investigation; Yan Xia was responsible for investigation; Shubin Li was responsible for data interpretation; Liyao Wang was responsible for supervision; Yuhao Chen was responsible for data acquisition; Yitian Du was responsible for data acquisition; Xinyu Li was responsible for supervision and substantively revised the manuscript.

## Data Availability Statement

The data supporting the findings and all plasmids and proteins are available upon reasonable request from the corresponding author.

