## Supplemental figures for "Entanglement-governed protein networks enable mechanically adaptive artificial skin for transplantation-scale skin replacement"

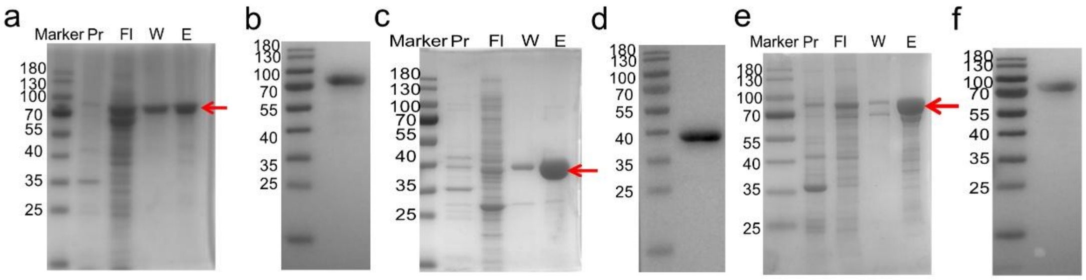


**Figure S1: Expression, purification, and validation of recombinant proteins.**
(a, c, e) SDS–PAGE analysis of purified FL4, FL8, and theFL8-Y→A mutant proteins. Protein samples were resolved on 12% separating gels and visualized by Coomassie Brilliant Blue staining. Lane M: molecular weight marker; lane Pr: pellet after cell lysis; lane Fl: flow-through fraction after $Ni^{2+}$ affinity chromatography; lane W: wash fractions containing low concentrations of imidazole; lane E: elution fractions containing high concentrations of imidazole. All target proteins migrated as single, well-defined bands at their expected molecular weights (FL4: 37 kDa;FL8: 76 kDa;FL8-Y→A: 76 kDa), indicating high purity.(b, d, f) Western blot analysis confirming the identity of the recombinant proteins. Immunoblotting was performed on the same samples as in (a) using an anti-His tag antibody. All proteins exhibited specific immunoreactive bands at the expected molecular weights, confirming the successful expression and purification of the target proteins.

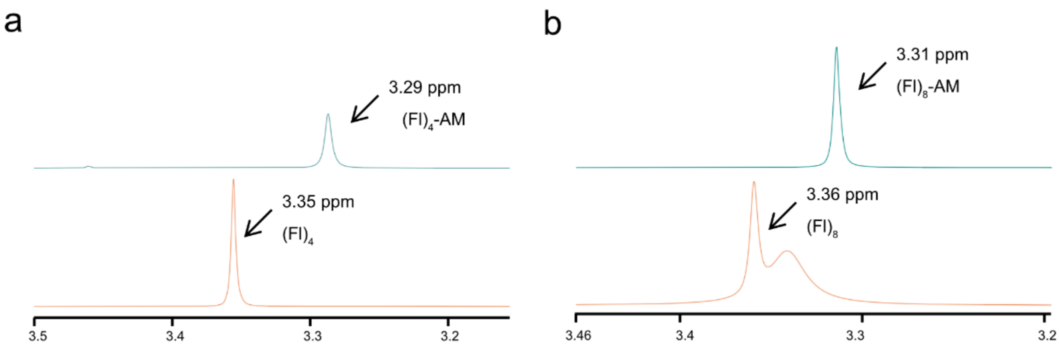


**Figure S2: ¹H NMR characterization of NHS-PEG-AM modification on FL4 and FL8 proteins.**

(a) Comparison of the ¹H NMR spectra of FL4 before and after modification. Lower panel: unmodified FL4 protein; upper panel: NHS-PEG-AM–modified FL4-AM protein. Upon modification, the characteristic proton resonance shifted from 3.35 ppm to 3.29 ppm, indicating successful conjugation of NHS-PEG-AM to the primary amine groups of the protein. (b) Comparison of the ¹H NMR spectra of FL8 before and after modification. Lower panel: unmodified FL8 protein; upper panel: NHS-PEG-AM–modified FL8-AM protein. Following modification, the corresponding characteristic proton signal shifted from 3.36 ppm to 3.31 ppm, confirming the successful functionalization of the FL8 protein.

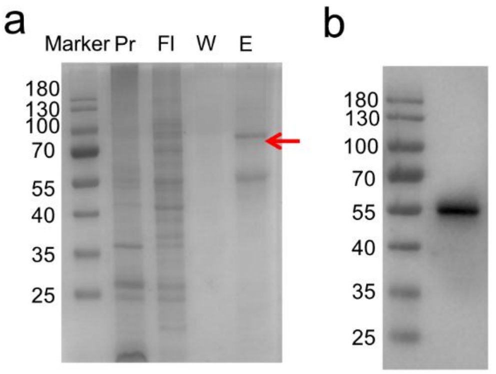


**Figure S3: Expression, purification, and validation of the recombinant CLP–EGF protein.**

(a) SDS–PAGE analysis of the purified CLP–EGF protein. Protein samples were resolved on a 12% separating gel and visualized by Coomassie Brilliant Blue staining. Lane M: molecular weight marker; lane Pr: pellet after cell lysis; lane Fl: flow-through fraction after $Ni^{2+}$ affinity chromatography; lane W: wash fractions containing low concentrations of imidazole; lane E: elution fractions containing high concentrations of imidazole. The target protein appeared as a single, well-defined band at the expected molecular weight, indicating high purity. (b) Western blot analysis confirming the identity of the recombinant protein. Immunoblotting was performed on the same samples as in (a) using an anti-His tag antibody. A specific immunoreactive band was detected at the expected molecular weight, confirming the successful expression and purification of the target protein.

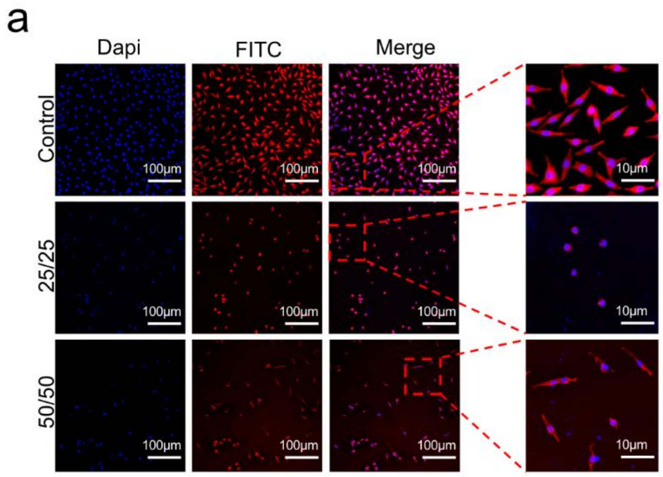


**Figure S4: Cytoskeletal spreading morphology of L929 cells on FED surfaces.** Phalloidin staining was used to visualize the cytoskeletal organization of L929 cells cultured on FED substrates. After 24 h of seeding, cells were stained with phalloidin (F-actin, red) and DAPI (nuclei, blue). Scale bar = 100 μm. The 25/25 group refers to cells cultured on FED surfaces composed of FL8-PEG-AM and AM at concentrations of 25 mg/mL each, whereas the 50/50 group corresponds to FED surfaces with both components at 50 mg/mL. Cells in the 50/50 group exhibited enhanced spreading and a typical spindle-shaped fibroblastic morphology.

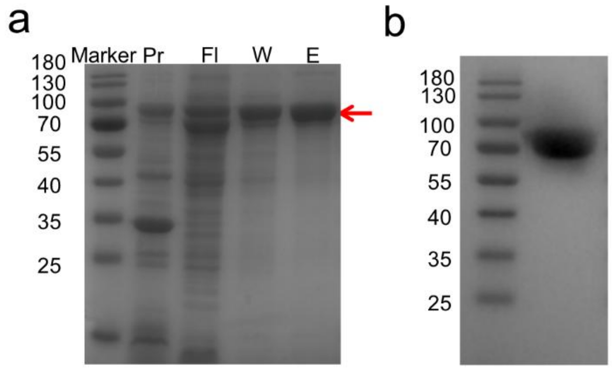


**Figure S5: Expression, purification, and validation of the recombinant FL8–ZINC protein.** (A) SDS–PAGE analysis of the purified FL8–ZINC protein. Protein samples were resolved on a 12% separating gel and visualized by Coomassie Brilliant Blue staining. Lane M: molecular weight marker; lane Pr: pellet after cell lysis; lane Fl: flow-through fraction after $Ni^{2+}$ affinity chromatography; lane W: wash fractions containing low concentrations of imidazole; lane E: elution fractions containing high concentrations of imidazole. The target protein appeared as a single, well-defined band at the expected molecular weight (~76 kDa), indicating high purity.(B)

Western blot analysis confirming the identity of the recombinant protein. Immunoblotting was performed on the same samples as in (A) using an anti-His tag antibody. A specific immunoreactive band was detected at the expected molecular weight, confirming the successful expression and purification of the target protein.

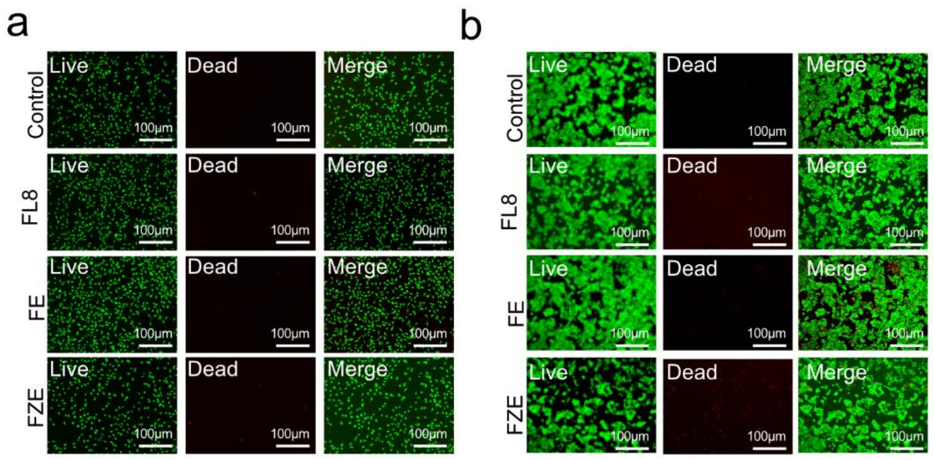


**Figure S6: Representative images of live/dead cell staining.**

L929 cells (a) and HaCaT cells (b) were co-cultured with extracts of different materials for 24 h, followed by staining with Calcein-AM (live cells, green) and propidium iodide (dead cells, red). Scale bar = 100 μm. The results show an increased number of dead cells in the FZE group, consistent with the CCK-8 assay results.

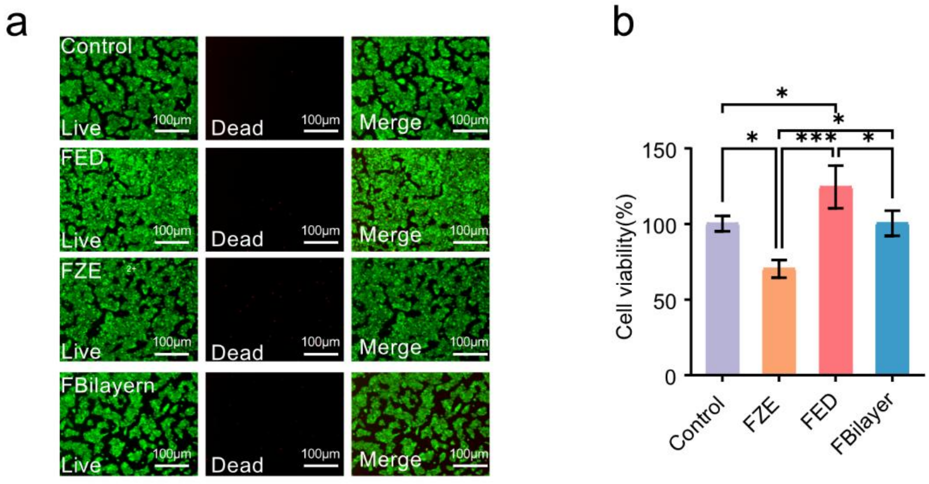


**Figure S7: Live/dead staining and CCK-8 assays for evaluating cytocompatibility of different material groups.**

(a) HaCaT cells were co-cultured with each material for 24 h, followed by staining with Calcein-AM (live cells, green) and propidium iodide (dead cells, red). Scale bar = 100 μm. The FBilayer group exhibited an intermediate level of cell viability, falling between the highly cytocompatible FED group and the moderately cytotoxic FZE group. (b) Cell viability was further quantified using the CCK-8 assay. HaCaT cells were incubated with extracts from each material group for 24 h prior to measurement. The FBilayer group consistently showed intermediate cell viability at all time points, compared with the FED and FZE groups. Data are presented as mean ± standard deviation (n = 5), with *$p < 0.05$ and **$p < 0.01$.

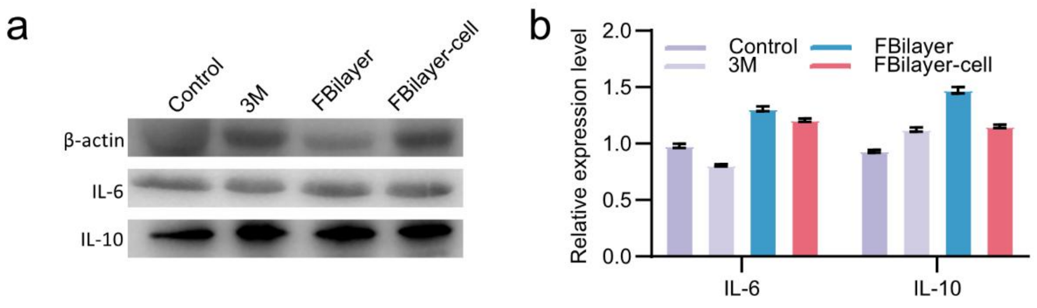


**Figure S8: Analysis of inflammatory cytokine expression (IL-6 and IL-10) in a murine large-area skin defect model.**
(a) Western blot analysis of IL-6 and IL-10 protein expression in wound-edge tissues at day 21 post-surgery. Samples were collected from the saline control group, the commercial dressing group (3M Tegaderm™), the FBilayer group, and the FBilayer + cell group. The expression levels of IL-6 (pro-inflammatory) and IL-10 (anti-inflammatory) were assessed, with β-actin used as the internal loading control. (b) Densitometric quantification of protein expression. Band intensities in (a) were analyzed using ImageJ software and normalized to β-actin. Quantitative results indicated no statistically significant differences in IL-6 or IL-10 expression among the treatment groups. Data are presented as mean ± standard deviation (n = 3), with n.s. indicating no significant difference.

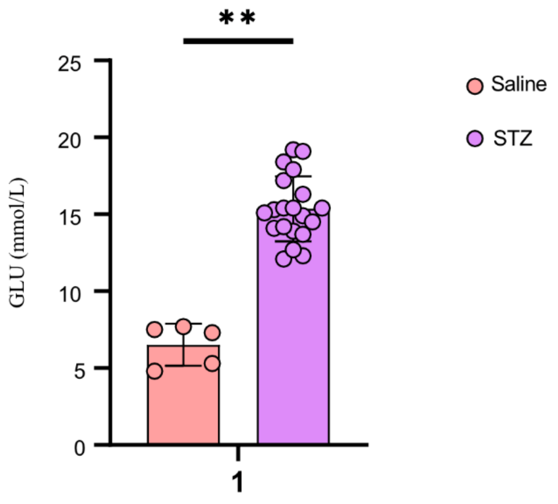


**Figure S9: Blood glucose levels in saline-treated and STZ-induced diabetic mice.** Blood glucose levels were measured following STZ administration to verify successful establishment of the diabetic model. Data are presented as mean ± s.d.; each dot represents one individual mouse. **$P$ < 0.01 versus the saline group.

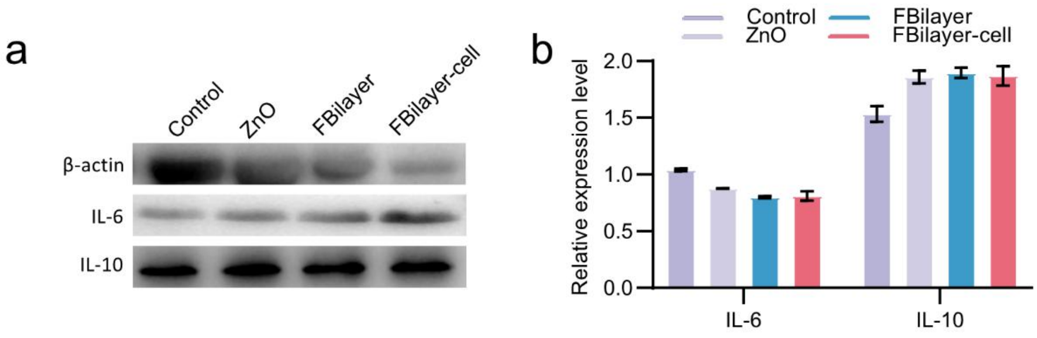


**Figure S10: Analysis of inflammatory cytokine expression (IL-6 and IL-10) in a diabetic mouse wound model.**

(a) Western blot analysis of IL-6 and IL-10 protein expression in wound-edge tissues of diabetic mice at day 12 post-surgery. Samples were collected from the saline control group, the ZnO ointment group, the FBilayer group, and the FBilayer + cell group. The expression levels of IL-6 (pro-inflammatory) and IL-10 (anti-inflammatory) were assessed, with GAPDH used as the internal loading control. (b) Densitometric quantification of protein expression. Band intensities in (A) were analyzed using ImageJ software and normalized to β-actin. Quantitative analysis showed no statistically significant differences in IL-6 or IL-10 expression among the treatment groups. Data are presented as mean ± standard deviation (n = 3), with n.s. indicating no significant difference.

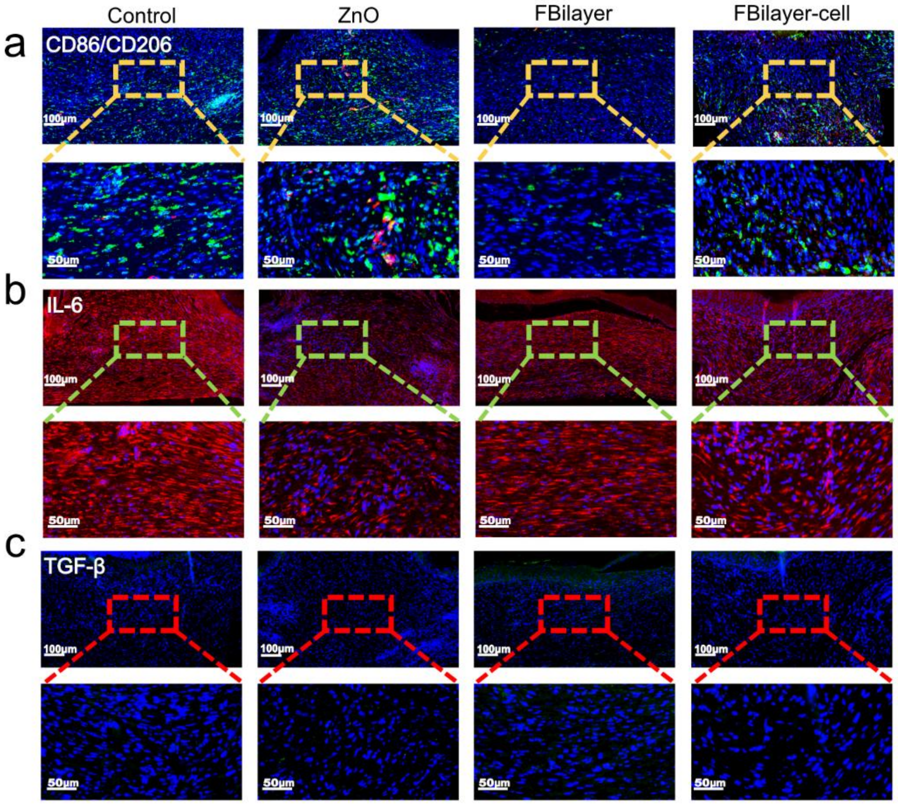


**Figure S11: Immunofluorescence analysis of macrophage polarization and inflammatory cytokine expression in a diabetic mouse wound model.**

(a) Representative immunofluorescence images showing macrophage polarization in wound tissues of diabetic mice at day 12 post-surgery. M1 macrophages were identified by CD86 staining (red), while M2 macrophages were labeled with CD206 (green); nuclei were counterstained with DAPI (blue). From left to right: saline control group, ZnO ointment group, FBilayer group, and FBilayer + cell group. Scale bar = 100 μm. Merged images indicate no obvious differences in the number or colocalization of CD86- and CD206-positive cells among the groups.(b, c) Immunofluorescence staining of inflammatory cytokines in wound tissues at day 12 post-surgery. The pro-inflammatory cytokine IL-6 (red) (b) and transforming growth factor-β (TGF-β, green) (c) were detected, with nuclei counterstained by DAPI (blue). From left to right: saline control group, ZnO ointment group, FBilayer group, and FBilayer + cell group. Scale bar = 100 μm. No significant differences in fluorescence intensity of IL-6 or TGF-β were observed among the groups.
